# Essential genes encoded by the mating-type locus of the human fungal pathogen *Cryptococcus neoformans*

**DOI:** 10.1101/2024.12.02.626420

**Authors:** Zhuyun Bian, Ziyan Xu, Anushka Peer, Yeseul Choi, Shelby J. Priest, Konstantina Akritidou, Ananya Dasgupta, Tim A. Dahlmann, Ulrich Kück, Minou Nowrousian, Matthew S. Sachs, Sheng Sun, Joseph Heitman

## Abstract

Fungal sexual reproduction is controlled by the mating-type (*MAT*) locus. In contrast to a majority of species in the phylum Basidiomycota that have tetrapolar mating-type systems, the opportunistic human pathogen *Cryptococcus neoformans* employs a bipolar mating-type system, with two mating types (**a** and α) determined by a single *MAT* locus that is unusually large (∼120 kb) and contains more than 20 genes. While several *MAT* genes are associated with mating and sexual development, others control conserved cellular processes (e.g. cargo transport and protein synthesis), of which five (*MYO2*, *PRT1*, *RPL22*, *RPL39*, and *RPO41*) have been hypothesized to be essential. In this study, through genetic analysis involving sporulation of heterozygous diploid deletion mutants, as well as in some cases construction and analyses of conditional expression alleles of these genes, we confirmed that with the exception of *MYO2*, both alleles of the other four *MAT* genes are indeed essential for cell viability. We further showed that while *MYO2* is not essential, its function is critical for infectious spore production, faithful cytokinesis, adaptation for growth at high temperature, and pathogenicity *in vivo*. Our results demonstrate the presence of essential genes in the *MAT* locus that are divergent between cells of opposite mating types. We discuss possible mechanisms to maintain functional alleles of these essential genes in a rapidly-evolving genomic region in the context of fungal sexual reproduction and mating-type evolution.

**IMPORTANCE:** Sexual reproduction is essential for long-term evolutionary success. Fungal cell type identity is governed by the *MAT* locus, which is typically rapidly evolving and highly divergent between different mating types. In this study, we show that the **a** and α alleles of four genes encoded in the *MAT* locus of the opportunistic human fungal pathogen *C. neoformans* are essential. We demonstrate that a fifth gene, *MYO2*, which had been predicted to be essential, is in fact dispensable for cell viability. However, a functional *MYO2* allele is important for cytokinesis and fungal pathogenicity. Our study highlights the need for careful genetic analyses in determining essential genes, which is complementary to high-throughput approaches. Additionally, the presence of essential genes in the *MAT* locus of *C. neoformans* provides insights into the function, maintenance, and evolution of these fast-evolving genomic regions.

## INTRODUCTION

Sexual reproduction is a fundamental process in the life cycle of eukaryotic organisms, playing a critical role in their long-term success. By reshuffling genetic material from two parents, sexual reproduction generates offspring with new combinations of traits and variable adaptive potential. This genetic diversity enables natural selection to act more effectively on populations, either by promoting the spread of beneficial mutations or by purging harmful mutations that have accumulated in parental genomes. Consequently, these processes enhance the population’s ability to adapt to environmental changes and improve long-term survival, highlighting the critical role of sexual reproduction in evolutionary success (1, 2).

In contrast to the X and Y chromosomes that determine sexual identity in humans, sexual reproduction in fungi is governed by less dimorphic chromosomal regions known as the mating-type (*MAT*) loci. Fungi typically employ one of two main mating-type systems: the bipolar and tetrapolar mating systems. In the Basidiomycota, mating type is generally determined by the tetrapolar mating system. This system involves two genetically and physically unlinked *MAT* loci: the *P/R* locus, which encodes the pheromones and pheromone receptor, and the *HD* locus, which encodes the transcription factors that govern sexual development. For sexual reproduction to occur, these two loci must differ between the mating partners (3, 4). Interestingly, members of the opportunistic human pathogenic *Cryptococcus* species complex, which belongs to the phylum Basidiomycota, instead have a bipolar mating system. In this system, the **a** and α mating types are determined by a single *MAT* locus carrying both the *P/R* and the *HD* genes (5, 6). *Cryptococcus* species, including *Cryptococcus neoformans*, can cause cryptococcal meningoencephalitis in both immunocompromised and immunocompetent individuals and result in more than 110,000 cryptococcal-related deaths annually (7–10).

Compare to the more compact *MAT* loci in ascomycetes, which only contain transcription factor genes, the *C. neoformans MAT* locus is unusually large (∼120 kb in size) and contains more than 20 genes (5). The *MAT***a** and *MAT*α alleles in *C. neoformans* exhibit considerable nucleotide divergence and extensive rearrangement, likely resulting from the lack of inter-allelic recombination (6, 11–15). In addition to genes that encode mating pheromones (*MF***a** or *MF*α), pheromone receptors (*STE3***a** or *STE3*α), and homeodomain transcription factors [*HD1* (*SXI1*α) or *HD2* (*SXI2***a**)] that are usually present in the two tetrapolar loci in this phylum, the *MAT* locus of *C. neoformans* also contains genes that are involved in mating (*STE11*, *STE12*, *STE20*), sporulation (*SPO14* and *RUM1*), and virulence (*CAP1*) (5, 16). Interestingly, five genes (*MYO2*, *PRT1*, *RPL22*, *RPL39*, and *RPO41*) encoded in the *C. neoformans MAT* locus have been predicted to be essential for viability (11). Of these, *MYO2* encodes a type V myosin motor protein whose ortholog in *Saccharomyces cerevisiae* is essential for mitochondrial inheritance (17), *PRT1* encodes a subunit of the eukaryotic translation initiation factor 3 (eIF3), *RPL22* and *RPL39* are two genes that encode ribosomal proteins that are important for translation, and *RPO41* encodes a mitochondrial RNA polymerase that is required for the transcription of mitochondrial genes.

Essential genes are crucial for the survival of an organism, making them potential drug targets for completely inhibiting the growth of pathogenic microbes, and research to identify these genes has been actively conducted. One common method involves identifying genes that cannot be deleted or disrupted; however, the possibility of transformation failure cannot be entirely excluded. Another widely used technique is high-throughput transposon mutagenesis sequencing (TN-seq), which has been applied to ascertain essential genes in fungi (18–21). However, it also comes with limitations that 1) results can be condition-specific, 2) different transposon systems may exhibit preferences for specific insertions sites, making it challenging to target genes uniformly across the genome, and 3) transposon insertions in one copy of an essential gene may not lead to loss of function in fungi with multiple genome or gene copies. An alternative approach involves deleting one copy of a gene in a diploid strain, inducing chromosome reduction to generate haploid progeny, and then demonstrating that a haploid mutant is inviable.

In this study, we assessed the essentiality of the five genes (*MYO2*, *PRT1*, *RPL22*, *RPL39* and *RPO41*) in the *C. neoformans MAT* locus by generating their heterozygous deletion mutants in the diploid *MAT***a**/α strain CnLC6683 using the transient CRISPR/Cas9 coupled with electroporation (TRACE) (22) technology, inducing sexual development and sporulation in these heterozygous deletion mutants, and then analyzing the phenotype as well as genotype of the resulting progeny. Our results demonstrated that, except for *MYO2*, all other alleles in this gene set are essential for viability. This result is consistent with a previous study confirming that *RPL22* and *RPL39* are essential by generating heterozygous deletion mutants in the AI187 diploid strain and analyzing the resulting progeny (23). Additionally, we validated the essentiality of these genes by employing regulatable promoters (a copper-regulated *CTR4* promoter or a Doxcycline-regulated Tet promoter) to control the expression of these genes. We then further investigated the function of Myo2 and found that both *myo2***a**Δ and *myo2*αΔ mutants exhibited defects in cytokinesis and displayed reduced vegetative fitness in a competition assay with the wild-type strains. Moreover, both **a** and α alleles of *MYO2* are important for vegetative growth at high temperature (37°C) and pathogenicity in the host. While the Myo2 ortholog in yeast plays an important role in mitochondrial inheritance (17), Myo2 was demonstrated not to be involved in mitochondrial uniparental inheritance in *C. neoformans*. In addition to the study of the *MYO2* gene, we generated and analyzed Ribo-seq and RNA-seq data from vegetative growth and mating samples of *RPL22* exchange allele strains to further study their possible role in sexual reproduction. Overall, this study confirmed the essentiality of four of the five predicted essential genes in the *MAT* locus. Further functional study of *MYO2* revealed its importance in cytokinesis, pathogenicity, and the production of infectious spores. We discuss our findings in the context of the origin, maintenance, and evolutionary trajectories of fast-evolving chromosomal regions such as the fungal *MAT* locus.

## RESULTS

### The *MAT* locus of *C. neoformans* encodes four essential genes

To study the essentiality of the genes within the *MAT* locus, we utilized a diploid strain CnLC6683(24), which was generated by fusing two congenic strains, KN99**a** and KN99α. Therefore, this diploid strain, CnLC6683, is homozygous throughout genome, except for the mating-type locus. Next, we deleted a single copy of each of the five genes, *MYO2*, *PRT1*, *RPL22*, *RPL39*, and *RPO41*, in the diploid strain CnLC6683 (Fig. 1A and 2A, see also Fig. S2 and S5C in the supplemental material). Because there are significant sequence divergence and rearrangements within *MAT*, we deleted the two opposite alleles (**a** with *NAT* and α with *NEO*) of each gene individually and generated ten heterozygous deletion mutants for the five predicted essential genes. Whole genome sequencing confirmed that all of these heterozygous deletion strains retained a diploid genome, and there were no segmental deletions linked to the gene deletions or in other genomic regions (Fig. S6). Phenotypic analyses of these heterozygous null mutants showed that, compared to the wildtype strain CnLC6693, they had similar vegetative fitness when grown on YPD solid medium. All of the heterozygous deletion strains exhibited robust hyphal growth and produced abundant basidiospores on MS medium. We did, however, observe a slight reduction in sporulation in the *PRT1***a**/*prt1*αΔ::*NEO* and *RPO41***a**/*rpo41*αΔ::*NEO* mutants (Fig. 1B, see also Fig. S3 in the supplemental material), In conclusion, our findings suggest a single allele of these genes in a hemizygous state is largely sufficient for mitosis and sexual reproduction.

**FIG 1.**
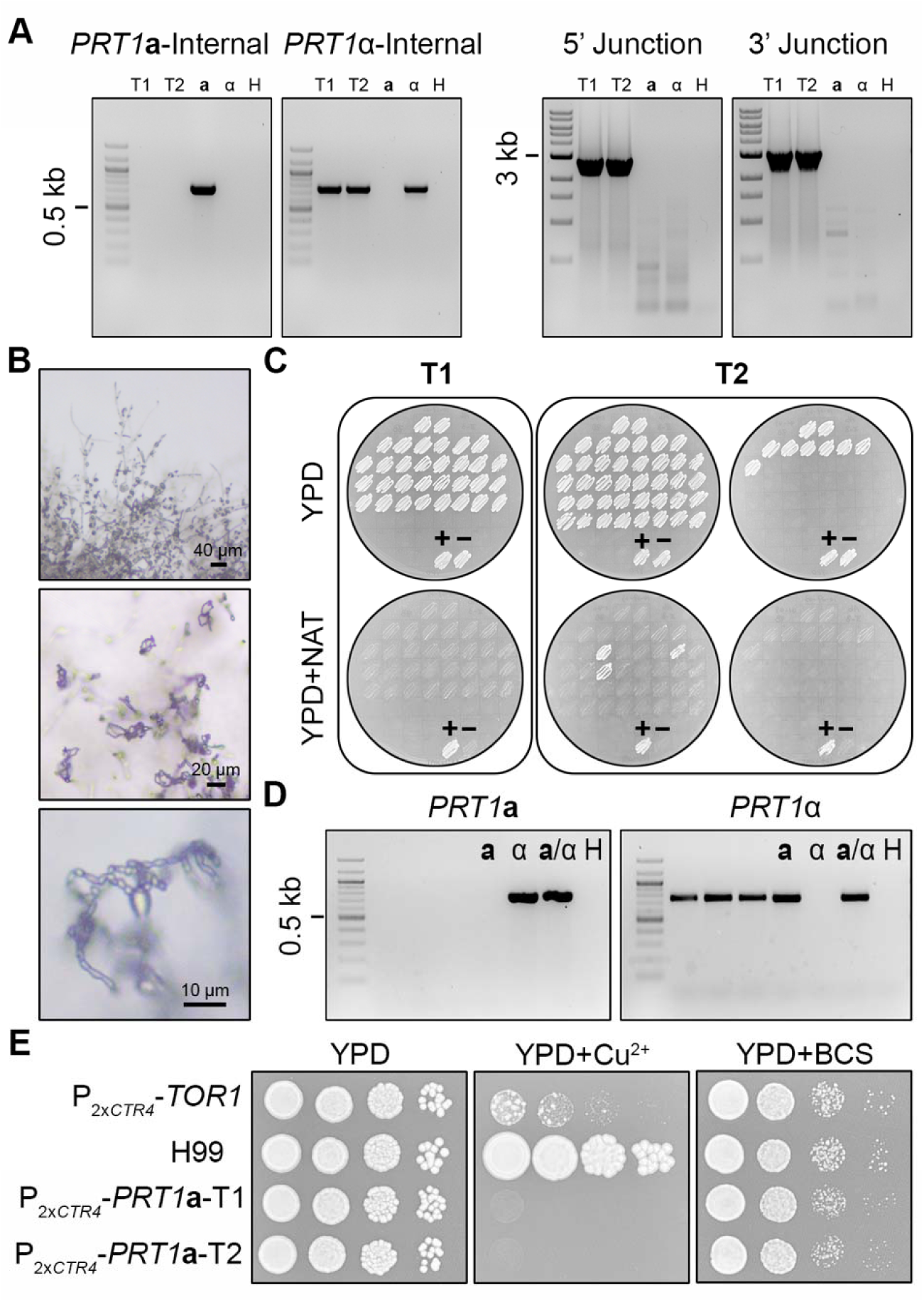
*PRT1***a** is an essential gene. (A) Genotype validation of *prt*1**a**Δ/*PRT*1α heterozygous deletion mutants with PCR targeting the internal regions of the ORFs of *PRT*1**a** and *PRT*1α (left), as well as the 5’ and 3’ junctions of the *prt*1aΔ::*NAT* allele. T1 and T2 are two independent transformants; **a**, α, and H indicate the KN99**a**, KN99α, and water controls for PCR, respectively. (B) The *prt*1aΔ/*PRT*1α heterozygous deletion mutants were wildtype for selfing and sporulation on MS media. (C) Phenotyping of germinated spores generated by two independent *prt*1**a**Δ/*PRT*1α mutants on YPD and YPD+NAT solid medium plates. The control (lower) patches are the parental diploid mutant strain as positive control (+) and wild-type strain CnLC6683 as negative control (−). (D) PCR with primer pairs targeting the internal regions of the ORFs of *PRT*1**a** and *PRT*1α confirmed the presence of a copy of wildtype *PRT*1α allele in the three *prt*1aΔ::*NAT* progeny, indicating of aneuploidy and consistent with *PRT*1 being essential for cell viability; **a**, α, and H indicate the KN99a, KN99α, and water controls for PCR, respectively. (E) WT, P*_2xCTR4_-*TOR1, and P*_2xCTR4_-*PRT1a strains were spotted and grown on YPD or YPD medium containing 200 μM BCS or 25 μM CuSO_4_. The plates were incubated at 30°C and photographed at 2 days after inoculation.

**FIG 2.**
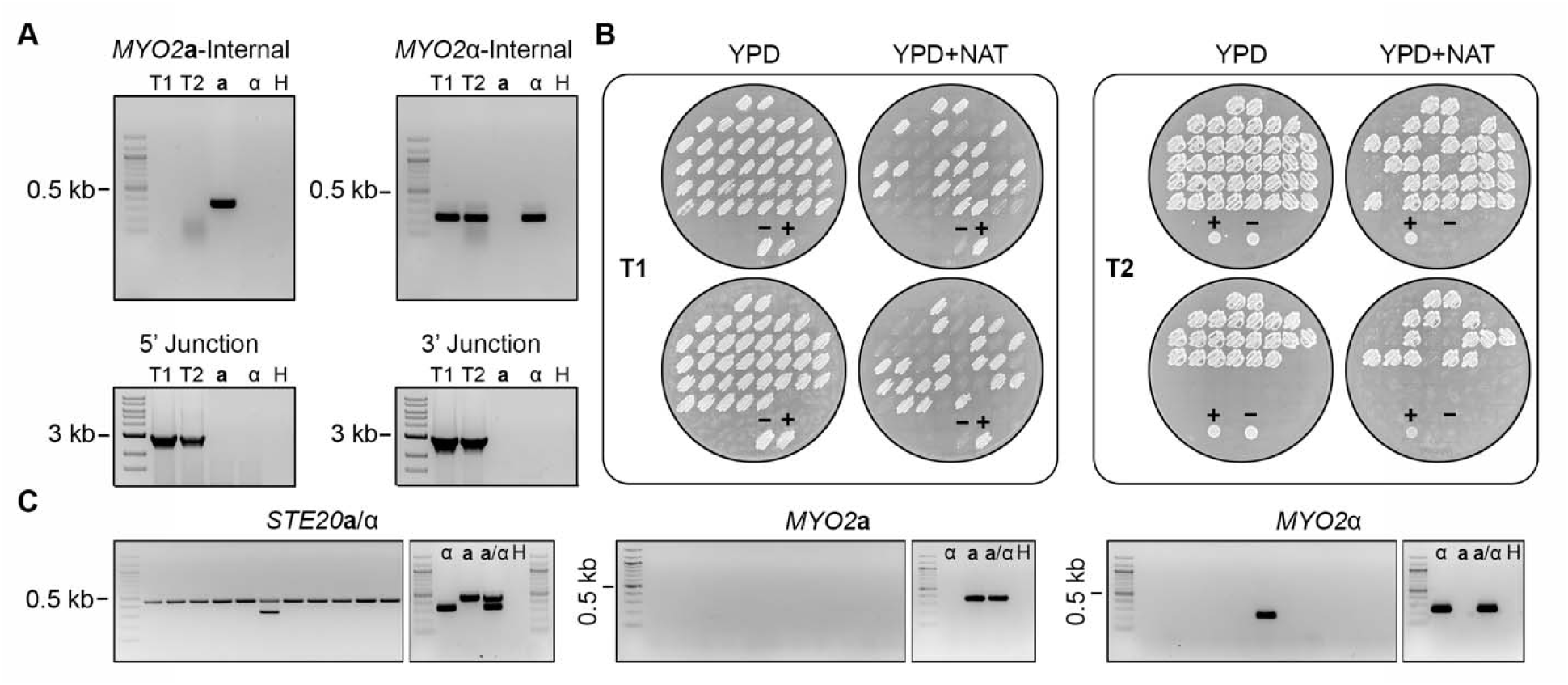
*MYO2***a** is not essential. (A) Genotyping of *myo2***a**Δ/*MYO2*α heterozygous deletion mutants with PCR targeting the internal regions of the ORFs of *MYO2***a** and *MYO2*α (left), as well as the 5’ and 3’ junctions of the *myo2***a**Δ::*NAT* allele. T1 and T2 are two independent transformants; **a**, α, and H indicate the KN99a, KN99α, and water controls for PCR, respectively. (B) Phenotyping of the germinated spores generated by two independent *myo2*aΔ/*MYO2*α mutants on YPD and YPD+NAT solid medium plates. For each transformant, ∼50% of the germinated progeny were NAT resistant. The control (lower) patches are the parental diploid mutant strain as positive control (+) and wild-type strain CnLC6683 as negative control (−). (C) PCR genotyping of a representative set of NAT resistant progeny with mating type specific primers targeting *STE20* (a and α), *MYO2***a**, and *MYO2*α, respectively, demonstrating that the vast majority of the NAT resistant progeny possessed neither MYO2a nor MYO2α, consistent with the gene being non-essential for viability; **a**, α, and H indicate the KN99**a**, KN99α, and water controls for PCR, respectively.

Each of the heterozygous deletion strains (e.g. *prt1***a**Δ::*NAT*/*PRT1*α) was then induced to undergo selfing, random haploid meiotic basidiospores were dissected, and drug resistance phenotype and genotype were analyzed. Our rationale is that if the gene is essential, then there should be no viable haploid meiotic progeny that inherit only the *MAT* allele containing the gene deletion mutation.

For each heterozygous deletion strain, we collected a minimum of 70 random meiotic basidiospores by microdissection, with germination rates ranging between 21% and 88% (Table 1). Phenotypic analyses showed that the vast majority of these viable progeny were sensitive to NAT (from those with deletions of the *MAT***a** allele) or NEO (from those with deletions of the *MAT*α allele) (Fig. 1C, see also Fig. S4A in the supplemental material). The heterozygous deletion strains producing the highest proportion of drug-resistant progeny were *MYO2***a**/*myo2*αΔ::*NEO* and the two independent *myo2***a**Δ::*NAT*/*MYO2*α strains (Fig. 2B, see also Fig. S5A and S5D in the supplemental material). Genotyping of these NAT/NEO resistant progeny from the *MYO2*/*myo2* heterozygous deletion strains showed that most do not possess the *MYO2***a** or *MYO2*α gene (Fig. 2C, see also Fig. S5B and 5E in the supplemental material), providing strong evidence that the *MYO2* gene is not essential. In contrast, except for five drug resistant progeny produced by *prt1***a**Δ::*NAT*/*PRT1*α (Fig. 1C), *rpl39***a**Δ::*NAT*/*RPL39*α and *rpo41***a**Δ::*NAT*/*RPO41*α (Fig. S4), other progeny that were randomly dissected from the heterozygous deletion strains of *PRT1*, *RPL22*, *RPL39*, and *RPO41* were all drug susceptible. Genotyping of the few drug-resistant progeny showed that all five still possessed a copy of the wildtype allele of the gene being deleted, but of the opposite mating type (Fig. 1D, see also Fig. S4B in the supplemental material). This is consistent with these progeny being aneuploid for chromosome 5, on which the *MAT* locus is located; it is also consistent with these genes being essential, in that the haploid progeny could inherit the deletion allele only if a wildtype allele (the opposite mating type allele in this case) was inherited simultaneously.

**Table 1.** Summary of spore viability analyses.

|  |  | Spores<br>dissected | Spores<br>germinated | Germination<br>rate | Number of<br>Drug<sup>R</sup> progeny | Presence of wild type allele in the<br>Drug<sup>R</sup> progeny <sup>a</sup> |
| --- | --- | --- | --- | --- | --- | --- |
| CnLC6683 |  | 70 | 62 | 89% | N/A |  |
| *prt1aΔ/PRT1α* | T1 | 96 | 32 | 33% | 0 |  |
|  | T2 | 96 | 49 | 51% | 3 | 100% |
| *rpl22aΔ/RPL22α* | T1 | 96 | 36 | 38% | 0 |  |
|  | T2 | 70 | 24 | 34% | 0 |  |
| *rpl39aΔ/RPL39α* | T1 | 96 | 43 | 45% | 0 |  |
|  | T2 | 70 | 30 | 43% | 1 | 100% |
| *rpo41aΔ/RPO41α* | T1 | 96 | 41 | 43% | 0 |  |
|  | T2 | 96 | 44 | 46% | 1 | 100% |
| *myo2aΔ/MYO2α* | T1 | 96 | 77 | 80% | 40 | 5% |
|  | T2 | 70 | 62 | 88% | 46 | 0% |
| *PRT1a/prt1αΔ* | T1 | 70 | 24 | 34% | 0 |  |
|  | T2 | 70 | 24 | 34% | 0 |  |
| *RPL22a/rpl22αΔ* | T1 | 140 | 63 | 45% | 0 |  |
|  | T2 | 70 | 25 | 36% | 0 |  |
| *RPL39a/rpl39αΔ* | T1 | 76 | 30 | 40% | 0 |  |
|  | T2 | 70 | 27 | 38% | 0 |  |
| *RPO41a/rpo41αΔ* | T1 | 70 | 42 | 60% | 0 |  |
|  | T2 | 96 | 40 | 41% | 0 |  |
| *MYO2a/myo2αΔ* | T1 | 250 | 53 | 21% | 29 | 6.8% |

<sup>a</sup> Only shown for the drug resistant colonies.

In addition to dissecting spores from heterozygous deletion mutants and then performing phenotypic and genotypic analysis of the meiotic progeny, we also took a different approach to test the essentiality for these genes. We inserted two tandem copper-regulated *CTR4* promoter (2x*CTR4*) upstream of the start codon of the *PRT1***a** gene (Fig. S2C) and then tested the viability on YPD medium supplemented with either copper sulfate (*CTR4*-repressing), or the copper chelator bathocuproine disulfonate (BCS, *CTR4*-inducing). The P_2x*CTR4*_-*TOR1* strain (25) served as a positive control. As shown in Fig. 1E, two independently constructed P_2x*CTR4*_-*PRT1***a** strains exhibited highly reduced growth under *CTR4*-repressing conditions (25 μM CuSO_4_) but grew as well as the WT strain when *CTR4* promoter was induced (200 μM BCS). Taken together, our analyses demonstrated that of the five genes predicted to be essential, four of them, *PRT1*, *RPL22*, *RPL39*, and *RPO41*, are indeed essential, while the remaining gene, *MYO2*, is dispensable for cell viability.

### The non-essential *MYO2* gene is required for cytokinesis, growth at 37**°**C, and pathogenicity

As the *MYO2* gene is not essential, we next conducted a comprehensive analysis of the gene using both *myo2***a**Δ and *myo2*αΔ haploid progeny obtained from selfing of the heterozygous deletion strains. We found that compared to the haploid wildtype controls, both *myo2***a**Δ and *myo2*αΔ deletion mutants showed significant growth defects when grown on YPD solid medium at 37°C, but not at 30°C (Fig. 3E). Interestingly, when grown in liquid YPD at 30°C, the wild-type strain H99 produced cells that were uniform and round, while cells produced by both mutant strains formed clusters (Fig. 3A). Hoechst staining showed proper nuclear division and migration in both deletion strains, even among cells forming clusters (Fig. 3C). Calcofluor white staining demonstrated accumulation and thickening of the calcofluor signal, at the mother-daughter cell connection sites (Fig. 3B). Thus, our results suggest that both *myo2***a**Δ and *myo2*αΔ mutants are defective in cytokinesis. Consistent with this observation, both *myo2*Δ mutants showed reduced vegetative fitness when compared to the wildtype control strains in a competition assay in liquid YPD at 30°C (Fig. 3D). Because some of the cells from *myo2***a**Δ and *myo2*αΔ mutants form clusters, the CFU of *myo2***a**Δ and *myo2*αΔ mutants might be underestimated. However, the declining proportion of mutant cells in the competition assay still indicates reduced fitness of both *myo2***a**Δ and *myo2*αΔ mutants in the competition assay. Taken together, our results showed that while *myo2***a**Δ and *myo2*αΔ were viable, there were considerable fitness costs associated with either gene deletion.

**FIG 3.**
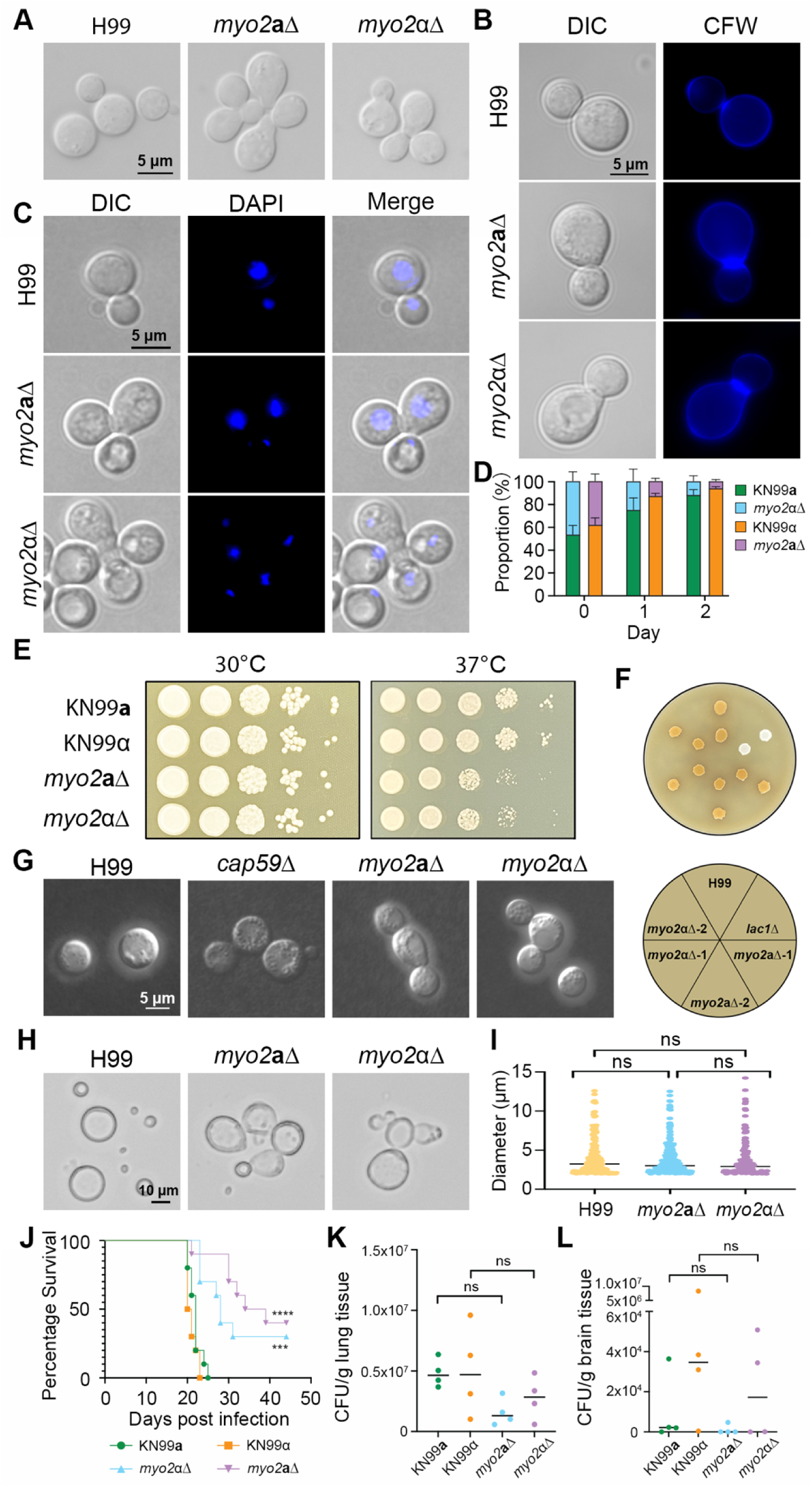
Phenotypic analyses of the haploid *myo2***a**Δ and *myo2*αΔ mutant progeny. (A) Both *myo2***a**Δ and *myo2*αΔ mutants exhibited compromised cytokinesis, manifested as cells forming abnormal clusters during vegetative growth in liquid YPD medium. Scale bar = 5 μm. (B and C) Microscopic images of cells from H99 wildtype, as well as *myo2***a**Δ and *myo2*αΔ mutants, after staining with Hoechst (C) or calcofluor white (B), confirming that both *myo2***a**Δ and *myo2*αΔ mutants exhibit normal nuclear division (C), but compromised cytokinesis (B) Scale bar = 5 μm. (D) Competition assay demonstrated that both *myo2***a**Δ and *myo2*αΔ mutants have reduced fitness compared to their respective wildtype strains, KN99a and KN99α. Plotted here are the percentages of the indicated mutants in co-cultures with their corresponding wildtype strains in liquid YPD medium after 0, 24, and 48 hours of incubation. Scale bar = 5 μm. (E) Compared to the wildtype strains, both *myo2***a**Δ and *myo2*αΔ mutant strains showed reduced growth on solid YPD medium at 37°C but not 30°C. (G) Cells stained with India ink showed no defects in capsule formation for either the *myo2***a**Δ or the *myo2*αΔ mutant strains. Scale bar = 5 μm. (F) No defects in melanin production were observed for *myo2***a**Δ or *myo2*αΔ mutant strains (H and I) Titan cells formed by *myo2***a**Δ and *myo2*αΔ mutant strains also showed compromised cytokinesis as they formed abnormal clusters (H), though no significant difference was observed in the proportion of titan cell between wild type and mutant strains (I). (J-L) Equal numbers of male and female A/J mice were infected intranasally with 10^6^ cells of the indicated WT and 1.5×10^6^ of *myo2***a**Δ and *myo2*αΔ mutant strains and analyzed for survival rate (n=10) and fungal burden (n=4).

We next investigated whether *MYO2* is involved in virulence and pathogenicity in *C. neoformans*. Virulence factors that have been identified in *C. neoformans* include the ability to grow at elevated temperature (37°C), production of an extracellular polysaccharide capsule, production of the cellular pigment melanin, and titan cell formation. While deletion of *MYO2* reduced vegetative fitness at 37°C (Fig. 3E), neither the *myo2***a**Δ nor the *myo2*αΔ mutant exhibited observable differences in the polysaccharide capsule thickness (Fig. 3F), melanin production (Fig. 3G), or titan cell formation when compared to the wildtype control strains (Fig. 3I), although compromised cytokinesis was observed in titan cells formed by both mutants (Fig. 3H).

We next examined the *in vivo* virulence of *myo2Δ* deletion strains in a murine inhalation infection model. We observed significantly prolonged survival in mice infected with either *myo2***a**Δ or *myo2*αΔ compared to the isogenic wildtype control (Fig. 3J); consistent with this, fungal burden analyses at 2-weeks post infection showed considerable, albeit not statistically significant, reduction in CFUs in both lungs and brains of mice inoculated with *myo2***a**Δ or *myo2*αΔ (Fig. 3K and L), when compared with their respective wildtype control. Taken together, our results suggest that *MYO2* plays an important role in virulence *in vivo*, which could be due to the reduced growth at 37°C observed in the deletion strains *in vivo*.

### *MYO2*a is required for sexual reproduction but dispensable for mitochondrial uniparental inheritance

In *C. neoformans*, mitochondria are uniparentally inherited (mito-UPI) from the *MAT***a** parent during **a**-α sexual reproduction (26). The ortholog of *MYO2* in *Saccharomyces cerevisiae* was demonstrated to be involved in mitochondrial inheritance (17). Thus, we sought to study whether *MYO2* is involved in mito-UPI during sexual reproduction in *C. neoformans*.

We analyzed three *myo2*Δ x wildtype unilateral crosses (Table 2). Two of these (crosses C1 and C2) were between the H99α wildtype strain and two independent *myo2***a**Δ mutant meiotic progeny that were each dissected from one of the two CnLC6683 *myo2***a**Δ::*NAT*/*MYO2*α heterozygous strains (Table 1). The third cross (cross C3) was between a *myo2*αΔ meiotic progeny dissected from the CnLC6683 *MYO2***a**/*myo2*α::*NEO* heterozygous strain (Table 1) and a strain in the KN99**a** background that possesses a mitochondrial genotype that is distinct from that of CnLC6683. Normal sexual development, including hyphal growth, basidia formation, and sporulation was observed in all three crosses, suggesting the presence of only one copy of the *MYO2* gene, either *MYO2***a** or *MYO2*α, is sufficient to complete sexual reproduction. Random basidiospores were dissected from both *myo2***a**Δ and *myo2*αΔ unilateral crosses (Table 2, crosses C1 to C3). The segregation of parental drug resistance markers among the progeny population from each cross showed high agreement with the expected frequencies for all of the phenotypic groups, suggesting there was no bias against any of the effected or of the predicted genotypes among the progeny, which was also consistent with the high spore germination rates observed in these crosses (Table 2). Genotyping of the mitochondria showed that of the more than 40 progeny analyzed for each cross, only one (from cross C3, Table 2; Fig. 4A and B) inherited the mitochondria from the *MAT*α parent. Thus, mito-UPI for the *MAT***a** parent is faithfully maintained during *myo2*Δ unilateral crosses.

**FIG 4.**
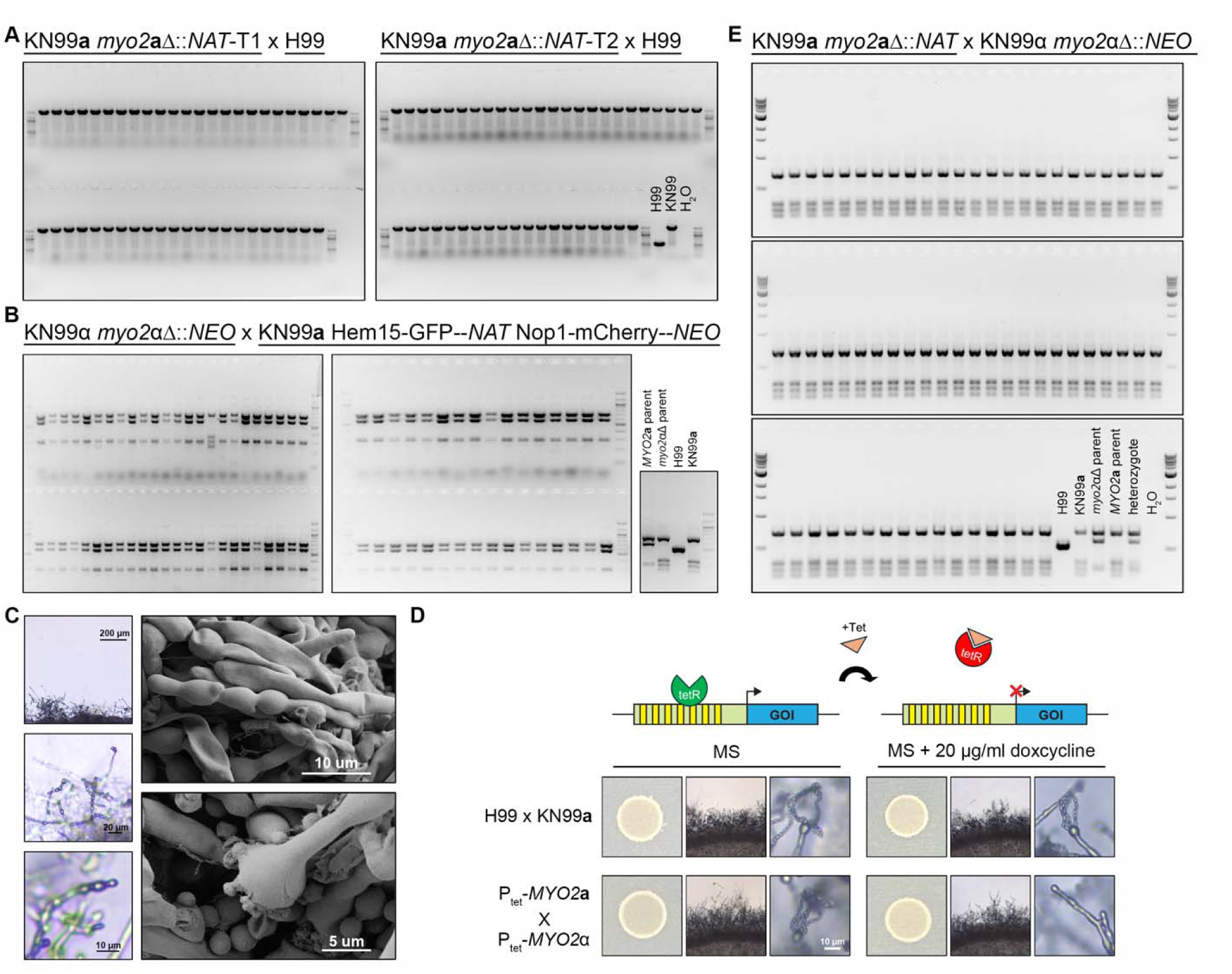
*MYO2***a** and *MYO2*α are not involved in mitochondrial uni-parental inheritance. (A-C) Genotyping of the mitochondrial types (COX1) of random spores dissected from *myo2***a**Δ and *myo2*αΔ unilateral (mutant x wildtype) (A and B) and bilateral (mutant x mutant) (E) crosses. (C) Bilateral cross of *myo2***a**Δ and *myo2*αΔ deletion strains showed defects in basidiospore formation. Scanning electron microscopy (SEM) analysis of hyphae and basidia from bilateral crosses showed hyphae with irregular segments and more than four budding sites on the basidial heads. Samples were prepared following incubation on MS media for one week. (D) Defects in basidiospore formation in bilateral cross of P_tet_-*MYO2***a** and P_tet_-*MYO2*α were observed on MS media containing 20 μg/ml doxycycline (right) but not on MS media (left).

**Table 2.** Summary of mito-UPI analyses of crosses involving myo2 deletion mutants.

| Cross type | Spores dissected | Spores germinated | Germination rate | No. of progeny analyzed | Number of progeny with *MATa* mitochondria genotype | Phenotypic group | Number of drug<sup>R</sup> progeny (%) | Expected frequency (%) |
| --- | --- | --- | --- | --- | --- | --- | --- | --- |
| C1 <sup>a</sup> | 56 | 46 | 82% | 46 | 46 (100%) | *NAT*<sup>R</sup> | 25 (54.3%) | 50% |
| C2 <sup>b</sup> | 56 | 43 | 77% | 43 | 43 (100%) | *NAT*<sup>R</sup> | 23 (53.4%) | 50% |
| C3 <sup>c</sup> | 92 | 80 | 87% | 48 | 47 (98%) | *NAT*<sup>R</sup> *NEO*<sup>R</sup> | 20 (41.7%) | 37.5% |
|  |  |  |  |  |  | *NAT*<sup>R</sup> *NEO*<sup>S</sup> | 4 (8.3%) | 12.5% |
|  |  |  |  |  |  | *NAT*<sup>S</sup> *NEO*<sup>R</sup> | 18 (37.5%) | 37.5% |
|  |  |  |  |  |  | *NAT*<sup>S</sup> *NEO*<sup>S</sup> | 6 (12.5%) | 12.5% |
| C4 <sup>d</sup> | 215 | 191 | 89% | 64 | 64 (100%) | *NAT*<sup>R</sup> | 31 (48%) | 50% |
|  |  |  |  |  |  | *NEO*<sup>R</sup> | 33 (52%) | 50% |

<sup>a</sup> KN99a *myo2aΔ::NAT*-T1 cross with H99.
<sup>b</sup> KN99a *myo2aΔ::NAT*-T2 cross with H99.
<sup>c</sup> KN99α *myo2αΔ::NEO* cross with KN99a Hem15-GFP--*NAT* Nop1-mCherry--*NEO*.
<sup>d</sup> KN99a *myo2aΔ::NAT* cross with KN99α *myo2αΔ::NEO*.

Interestingly, when we set up *myo2***a**Δ x *myo2*αΔ bilateral mutant crosses for mito-UPI analyses, we observed several defects in sexual development, including significantly impaired basidium formation and sporulation, with distended segments along the hypha (Fig. 4C). This suggests that sexual development and sporulation are highly compromised when both copies of *MYO2* are absent. To confirm this, we engineered haploid *MAT***a** and *MAT*α strains in which their respective *MYO2***a** and *MYO2*α genes were under the Tet-off regulatable promoter, where the expression of the gene can be repressed by the presence of exogenous doxycycline in the growth medium (Fig. 4D) (27). While the cross between the Tet-*MYO2***a** and Tet-*MYO2*α strains appeared to be normal and indistinguishable from the wildtype cross between H99α and KN99**a** on MS medium without doxycycline, on MS medium supplemented with doxycycline (20 μg/ml), the same cross exhibited impaired sexual development similar to that observed in *myo2*αΔ x *myo2***a**Δ bilateral crosses. We further showed that the defect in sexual development was not due to the mere presence of doxycycline in the medium, as the crosses between wildtype strains H99α and KN99**a** appeared to be identical on MS and MS+doxycycline media (Fig. 4D). Thus, our results strongly suggest that a functional *MYO2* gene, from either parent, is indispensable for successful and complete sexual development and production of infectious spores.

Due to the severe sporulation defects observed in the *myo2*αΔ x *myo2***a**Δ bilateral crosses, we opted instead to dissect and analyze blastospores, which are yeast cells that bud off hyphae, for the analyses of mitochondrial inheritance. Because previous studies have shown that, in *C. neoformans*, mito-UPI is established during zygote formation and completion by early stages of hyphal development (26), we reasoned that the mitochondrial type of the blastospores budding from the hyphae should be identical to the type in the hyphae, as well the type in the eventual basidiospores. We observed similarly high germination rate in the blastospores, with 1:1 segregation of the two parental drug markers, which again suggests there was no underrepresentation of any genotypic groups (Table 2, cross C4). Genotyping of the mitochondrial genome showed that all 64 blastospores analyzed inherited mitochondria from the *MAT***a** parent (Fig. 4E), suggesting mito-UPI is also maintained in *myo2*αΔ x *myo2***a**Δ bilateral crosses.

Taken together, our results showed that the *MYO2* gene is critical for robust sexual reproduction and sporulation in *C. neoformans*; however, it is not required for uniparental inheritance of mitochondria.

### Rpl22 modulates translation dynamics during sexual reproduction

The *RPL22* gene encodes ribosomal protein L22, a component of the 60S large ribosomal subunit. The **a** and α alleles differ by five amino acids that are located close to the N-terminus. In our previous studies, we generated two Rpl22 allele-exchange strains: YFF96α (*rpl22*α::*RPL22***a**) that is isogenic to YFF92 (23), in which the *RPL22*α allele in strain H99α was replaced with the *RPL22***a** allele derived from strain KN99**a**; and YFF113**a** (*rpl22***a**::*RPL22*α^N^-*RPL22***a**^C^), in which the *RPL22***a** allele in strain KN99**a** was genetically modified to replace the **a**-specific amino acids at the N-terminus with their respective α-specific variants, and thus, this strain has a functional *RPL22*α allele (23). In this study, we conducted a series of ribosome profiling (Ribo-seq) and RNA-seq analyses of these two strains, both in solo-cultures as well as in crosses (unilateral and bilateral) and compared them to their corresponding wildtype background controls (Fig. 5).

**FIG 5.**
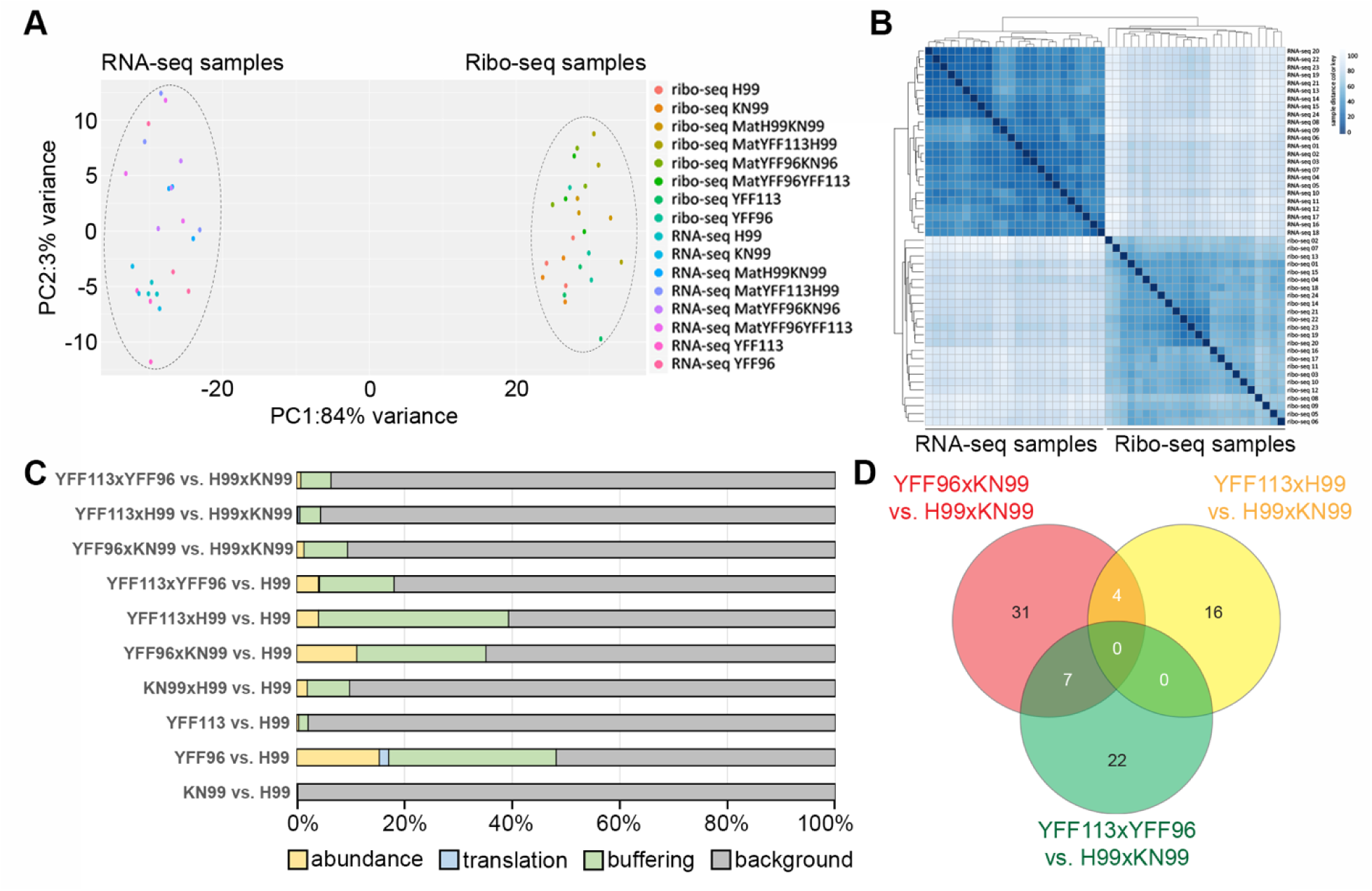
Analysis of RNA-seq and Ribo-seq data from *C. neoformans* mutant strains and genetic crosses. (A) Principal component analysis of samples after read counts were normalized with DESeq2. Ribo-seq samples group separately from RNA-seq samples. Limited correlation of RNA-seq and Ribo-seq results was previously observed in studies with other systems and does not in itself preclude drawing conclusions from the data (28–30). (B) Heatmap of sample distances of read counts normalized with DESeq2. Similar to the principal component analysis, Ribo-seq samples cluster separately from RNA-seq samples. (C) Bar chart showing the percentage of genes in the four categories for each sample condition. Only genes (total of 520) that categorized to the abundance, translation or buffering group in at least one of the comparisons were included. (D) Venn diagram of genes in the category buffering in comparisons of mating samples versus mating of strains KN99 and H99. Numbers of genes that are in the category buffering in one or more comparisons based on anota3seq are indicated.

Similar to what was observed in studies with other systems, the RNA-seq and Ribo-seq cluster separately (Fig. 5A and B), which in itself does not preclude drawing conclusions from the data (28–30). Based on the transcription and translation levels, we further classified the transcription/translation profiles of the genes into four categories: 1) changes in mRNA abundance (i.e. “abundance”, referring to proportional significant changes in both total mRNA and translated mRNA), 2) changes in translational efficiency (i.e. “translation”, referring to significantly disproportionate changes in translation levels relative to their total mRNA levels), 3) buffering (referring to stable translation levels even when significant changes were observed in the mRNA levels), and 4) background (referring to genes for which no significant changes were observed for translation and mRNA levels). A total of 520 genes were categorized to the abundance, translation, or buffering group in at least one of the comparisons and used for further analysis (Fig. 5C).

Overall, we observed that most of the genes did not show changes in their expression profiles compared to the controls (i.e. the “background” category). Among the genes that showed significant differences, the majority of them belonged to the “buffering” and “abundance” categories, and very few belonged to the “translation” category, suggesting the changes in the expression profiles were usually associated with changes in gene transcription.

In solo cultures, while both KN99**a** and the *RPL22* exchanged strain YFF133**a** showed minimal differences when compared to the H99α control strain, the *RPL22* exchanged strain YFF96α exhibited considerable changes, with 80, 9 and 162 genes that were differentially regulated at the level of abundance, translation, and buffering, respectively. This asymmetrical effect observed between YFF133**a** and YFF96α indicates that the expression of *RPL22***a** in a *MAT*α background led to more significant changes in gene transcription and translation compared to the reciprocal expression of *RPL22*α in the *MAT***a** background (Fig. 5C).

For the samples from crosses involving YFF96α and YFF133**a** (unilateral) or both (bilateral), all of them showed clear differences in transcription and translation compared to wildtype controls, and overall more changes were identified when compared to H99α solo culture than when crosses between H99α and KN99**a** were used as controls, consistent with metabolic changes occurring during the physiological transition from vegetative yeast growth to sexual development (Fig. 5C). Specifically, when compared to mating of strains KN99**a** and H99α, crosses involving YFF96α and YFF133**a** (two unilateral crosses) or both (one bilateral cross) had 42, 20 and 29 genes that were differentially regulated at the level of buffering, respectively (Fig. 5D). Interestingly, there was no gene that was found to be similarly differentially regulated in all three crosses. Additionally, only a small number of differentially regulated genes were found to be shared between crosses YFF96α x KN99**a** and YFF113**a** x H99α, or between YFF96α x KN99**a** and YFF113**a** x YFF96α; no common differentially regulated gene were found between crosses YFF113**a** x H99α and YFF113**a** x YFF96α (Fig. 5D). This suggests that the **a** and α alleles of *RPL22* might encode proteins that are regulated quite differently during sexual reproduction and changes in any allele could lead to changes in regulation of divergent sets of genes.

## DISCUSSION

Five genes (*MYO2*, *PRT1*, *RPL22*, *RPL39*, and *RPO41*) in the *MAT* locus of *C. neoformans* were predicted to encode proteins required for viability (11). In our study, we confirmed that all but one (*MYO2*) were indeed essential. *MYO2* is not essential its deletion led to severe defects in cytokinesis. While the DNA sequence can be highly divergent between the **a** and α alleles of these genes, their predicted protein structures are highly similar (Fig. S1A), reflecting their importance. Our results are also consistent with the predictions of essentiality based on a high-throughput transposon mutagenesis and sequencing system (Tn-Seq) in *C. neoformans* (21) (Fig. S1B). Additionally, two of the genes, *RPL22* and *RPL39*, had been previously predicted to be essential in *C. neoformans* based on the analyses using a heterozygous diploid strain, AI187, that was derived from the fusion of two haploid strains, JF99 (*MAT***a** *ura5*) and M001 (*MAT*α *ade2*), the latter of which had undergone random UV mutagenesis to generate the *ade2* mutant together with ∼200 other extraneous mutations (31). In contrast to AI187, strain CnLC6683 (24) is a fusion product of the congenic strain pair KN99**a** and KN99α. It is fully prototrophic and does not have the *ade2* and *ura5* auxotrophic mutations nor the random mutations in the AI187 genome that were introduced by the M001 genome, which could in some cases complicate genetic analysis. Thus, our study utilizing the strain CnLC6683 presents the most definitive evidence for the essentiality of these genes.

We employed a doxycycline regulatable promoter to modulate the expression of *MYO2***a** and *MYO2*α to examine their roles more closely. On MS solid medium supplemented with 25 μM doxycycline, defects in spore production could be observed in bilateral cross between P_tet_-*MYO2***a** and P_tet_-*MYO2*α, indicating that the Tet-off system worked as expected for the non-essential genes *MYO2***a** and *MYO2*α. However, we failed in generating mutants with a doxycycline regulatable promoter for the four essential genes. Neither Tet-off promoter integrated transformants could be obtained from transformation using haploid wild type strains, nor could drug-resistant progeny be recovered from sporulation of diploid heterozygous mutants, when a doxycycline regulatable promoter was integrated in front of essential genes. We then utilized the *CTR4* promoter for the regulation of these essential genes. Our findings showed that to achieve robust regulation, a tandemly duplicated *CTR4* promoter needed to be inserted between the endogenous promoter regions and the start codon of the genes, suggesting that the original promoter sequences are critical for the proper function of the essential genes, even when they have been placed under the control of an extraneous promoter system.

While *MYO2* is a non-essential gene in *C. neoformans*, its ortholog in *S. cerevisiae*, *ScMYO2*, is an essential gene. Because ScMyo2 was reported to play a major role in mitochondrial motility (17), we investigated whether CnMyo2 plays a role in the mitochondrial uniparental inheritance during sexual reproduction. No defects in mito-UPI was observed for either unilateral cross between *myo2***a**Δ or *myo2*αΔ mutants and wild-type strains or a bilateral cross between *myo2***a**Δ and *myo2*αΔ. However, defects in cytokinesis were observed in both *myo2***a**Δ and *myo2*αΔ mutants. The successful completion of cytokinesis in animal and fungal cells requires the involvement of actomyosin ring (AMR) contraction (32, 33). In *S. cerevisiae*, the type II myosin ScMyo1 was reported to be important for forming the ring (34) and ScMlc1 is a light chain for both ScMyo1 and the type V myosin ScMyo2 that coordinates AMR function, membrane trafficking, and septum formation during cytokinesis (35). It is possible that Myo2 in *C. neoformans* is also involved in AMR function as deletion of *MYO2* results in defects in cytokinesis.

We observed that while *PRT1***a**/*prt1*αΔ and *RPO41***a**/*rpo41*αΔ mutants exhibited normal hyphal development, they both showed reduced sporulation. Interestingly, normal sporulation was observed in their respective reciprocal deletion strains, *prt1***a**Δ/*PRT1*α and *rpo41***a**Δ/*RPO41*α, indicating the presence of asymmetrical requirements for the **a** and α alleles for faithful sexual development. This could be due to haploinsufficiency for robust sporulation of the **a** alleles, or mating-type specific activities or functions of the genes. Notably, *RPO41***a** and *PRO41*α share 99.3% identity in nucleotide sequence and 97.59% identity in protein sequence, with the main difference being a 23 amino acid region located at the C-terminus of *RPO41***a** that is absent in *RPO41*α. It will be interesting to know whether this short amino acid sequence causes functional and/or regulatory differences between the products of the *RPO41***a** and *RPO41*α alleles. Asymmetrical characteristics were also observed in RNA-seq and Ribo-seq analyses of Rpl22**a** and Rpl22α, where the expression of the *RPL22***a** allele in a *MAT*α background (YFF96α) induced significantly more transcription/translation changes compared to the reciprocal expression of the *RPL22*α allele in a *MAT***a** background (YFF113**a**). This is also consistent with previous studies that have shown that asymmetry is present in the expression of pheromone and pheromone receptors, as well as in the early sexual development and morphogenesis of **a** and α cells (10, 36).

An interesting question is how are these essential genes maintained in the *MAT* locus? One characteristic of *MAT* is the highly repressed recombination within this locus during meiosis (5, 37).This could help maintain mating locus specific alleles, although it also facilitates the accumulation of deleterious mutations and impedes their removal. Additional mechanisms could contribute to the maintenance of proper function of essential genes in the *MAT* locus. For example, gene conversion occurs within the *MAT* locus during **a**-α sexual reproduction (38).

Gene conversion can remove detrimental mutations by employing the opposite allele as a template, and consequently lead to slower evolutionary divergence between the two alleles. This is consistent with the observed sequence identity between the **a** and α alleles of the non-essential gene *MYO2* (58%), and essential genes *PRT1* (84%), *RPL22* (88%), *RPL39* (90%), and *RPO41* (99%). Additionally, recombination hot spots flanking the *MAT* locus could potentially facilitate the removal of the *MAT* allele containing deleterious mutations as a whole (39). Moreover, the mating-type locus is free to recombine during unisexual reproduction, facilitating the removal of potential deleterious mutations (37). The presence of essential genes could have contributed to the initial formation and maintenance of the unusually large and highly rearranged *MAT* locus, as ectopic recombination within *MAT* would likely result in recombinants that are missing essential genes, rendering them inviable (4, 13, 40). Essential genes have been found within the *MAT* loci in other fungal species. For example, two essential genes, *PIK* and *PAP*, have been identified in the *MAT* locus of *Candida albicans* (41), of which *PIK* encodes a phosphatidylinositol kinase involved in signal transduction (42), while *PAP* encodes a poly(A) polymerase that polymerizes the polyadenosine tail at the 3’ ends of mRNAs (43). Interestingly, while located in the mating-type locus, neither of these two genes have known functions related to mating (41, 44). It is possible that these essential genes are maintained in the mating-type locus as it could provide evolutionary advantages by imposing a selective pressure that maintains both mating capabilities and essential cellular functions within a diverging genomic region.

Our results further confirmed the presence of co-evolution of genes, as well as their regulatory sequences, within the **a** and α *MAT* alleles of *C. neoformans*, respectively. Further research, such as functional analyses of the essential genes utilizing conditional alleles, will further shed light on the formation, maintenance, and evolution of the *MAT* locus, as well as provide insight into other rearranged genomic regions, such as sex chromosomes.

## MATERIALS AND METHODS

### Ethics statement

All animal experiments in this manuscript were approved by the Duke University Institutional Animal Care and Use Committee (IACUC) (protocol #A098-22-05). Animal care and experiments were conducted according to IACUC ethical guidelines.

### Strains and culture conditions

Heterozygous mutants were generated in the diploid *C. neoformans* strain CnLC6683 (24). For transformation of haploid *C. neoformans* strains, we employed H99α and KN99**a** (45). All of the strains were maintained on YPD (1% yeast extract, 2% Bacto Peptone, 2% dextrose) agar medium. Mating and cell mass collected for Ribo-seq and RNA-seq were conducted on Murashige and Skoog (MS) (Sigma-Aldrich M5519) plates at incubate at room temperature in the dark.

### Construction of heterozygous deletion and promoter replacement strains

For generation of heterozygous mutants, the *NAT* or *NEO* gene expression cassette were amplified from plasmids pAI3 and pJAF1, respectively. Approximately 1.5 kb regions (homologous arms) flanking the genes of interest were amplified from H99α for *MAT*α alleles or KN99a for *MAT***a** alleles genomic DNA and fused with the *NAT* (*MAT***a** alleles) or *NEO* (*MAT*α alleles) drug resistance marker with overlapping PCR as previously described (46) to generate the doner DNA cassettes. CRISPR-Cas9-directed mutagenesis was used for the mutant generation. The *CAS9* cassette was PCR-amplified from plasmid pXL-1 with universal primers M13F and M13R (46). The desired target sequences for the sgRNA constructs were designed using the Eukaryotic Pathogen CRISPR guide RNA/DNA Design Tool (EuPaGDT) with default parameters (47). Two gRNAs were designed and used for each gene of interest. Complete gRNAs were generated by one-step overlap PCR as described previously (23). 1.5 μg donor DNA cassette, 400 ng *CAS9* cassette and 150 ng of each complete gRNAs fragment were mixed and condensed to a 5 μL volume before introduced to the diploid strain CnLC6683 with the transient CRISPR/Cas9 coupled with electroporation (TRACE) transformation approach (46).

To construct promoter replacement strains, KN99**a** strains were used. The native promoter of *PRT1***a** was replaced with the two tandem *CTR4* promoter amplified from P_2x*CTR4*_-*TOR1* strain (25) with primer JOHE54314/ZB363 and JOHE54314/ZB364 (Table S2). Similar strategy using TRACE transformation approach was applied to generate the mutants. To induce copper sufficiency or deficiency, YPD plates were supplemented with 25 μM CuSO_4_ or 200 μM of the copper chelator bathocuproine disulfonate (BCS).

To generate the doxycycline regulatable promoter strain for *MYO2***a** and *MYO2*α, ∼300 bp of the original promoter in front of their coding DNA sequencing was replaced with the Tet promoter that amplified from vector pCL1774 (27) with universal primer M13F/M13R (Table S2). The TRACE transformation approach (22) was applied to generate the mutants. 25 μM doxycycline was added to corresponding media to induce the Tet-off system.

### Whole genome sequencing and ploidy analysis

Illumina sequencing of the strains was performed at the Duke sequencing facility core (https://genome.duke.edu/), using Novaseq 6000 as 150 paired-end sequencing. The Illumina reads, thus obtained, were mapped to the H99 genome assembly using Geneious (RRID:SCR_010519) default mapper to estimate ploidy. The resulting BAM file was converted to a. tdf file, which was then visualized through IGV to estimate the ploidy based on read coverage for each chromosome.

### Self-filamentation, mating and genotyping

To analyze the essentiality of a target gene, a colony size of YPD culture of a generated heterozygous deletion mutant was resuspended in sterilize water and 4 μL was spotted onto Murashige and Skoog (MS) (Sigma-Aldrich M5519) plates. Inoculated MS plates were then incubated at room temperature in the dark for 10 days, and random spores were then dissected as previously described (48). Germinated individual spores were transferred and patched onto fresh YPD and YPD contain NAT or NEO medium, and genomic DNA of progeny that grown on YPD with drug plates was extracted from the biomass as described in a previous study (38).

To test the effect of a *MYO2***a** or *MYO2*α on mito-UPI, the unilateral and bilateral crosses were set up by spotting the mixture of the two parental strains onto MS medium, incubated at room temperature in the dark for 10 days, and random spores were then dissected, patched onto fresh YPD and YPD contain NAT or NEO medium and used for genomic DNA extraction as described above. The mitochondrial genotypes between H99 and KN99 were differentiated with PCR markers targeting the presence/absence of introns in the *COX1* gene, as previously described (49). For one of the unilateral crossings, *myo2*αΔ was crossed with KN99**a** that contains a recombinant mitochondrial genotype, which can be differentiated from a wild-type KN99 mitochondrial type by RFLP digestion with *Bsr*I. Therefore, mitochondrial genotyping for this crossing was based on PCR-RFLP markers targeting the *COX1*.

### Imaging with light microscopy and SEM

Brightfield and differential interference contrast (DIC) microscopy images were visualized with an AxioScop 2 fluorescence microscope and captured with an AxioCam MRm digital camera (Zeiss, Germany). Consistent exposure times were used for all images analyzed.

For sample preparation for SEM from self-filamenting diploid strains, an agar slice of the plated cells was fixed in a solution of 4% formaldehyde and 4% glutaraldehyde for 16 hours at 4°C. The fixed cells were then gradually dehydrated in a graded ethanol series (30%, 50%, 70%, and 95%), with a one-hour incubation at 4°C for each concentration. This was followed by three washes with 100% ethanol, each for 1 hour at room temperature. The samples were further dehydrated using a Ladd CPD3 Critical Point Dryer and coated with a layer of gold using a Denton Desk V Sputter Coater (Denton Vacuum, USA). Hyphae, basidia, and basidiospores were observed with a scanning electron microscope with an EDS detector (Apreo S, ThermoFisher, USA).

### Competition assay

KN99**a**, KN99α, *myo2***a**Δ and *myo2*αΔ strains were cultured overnight at 30°C in liquid YPD or YPD+NAT (*myo2***a**Δ)/NEO (*myo2*αΔ). Cells were adjusted to equal densities using OD_600_ measurements and mixed in equal numbers in a 4 mL YPD co-culture. KN99**a** is mixed with *myo2***a**Δ and KN99α is mixed with *myo2*αΔ. This plating process was repeated at 24 and 48 hours to calculate the cell density of each strain in the co-culture. The data presented are based on four biological replicates, each with three technical replicates.

### Melanin and capsule formation analysis and serial dilution assays

Fresh cells were spotted onto Niger seed (7% Niger seed, 0.1% dextrose) plates and incubated at 30°C for three days to assay the melanin formation. For capsule analysis, strains were incubated for 2 days in RPMI (Sigma-Aldrich R1383, 2% dextrose) liquid media at 37°C, followed by negative staining with India ink. To test the growth ability of *myo2* mutants at 37°C, fresh cells of KN99**a**, KN99α, *myo2***a**Δ and *myo2*αΔ were diluted to a starting OD_600_ of 1, serially diluted 10-fold, and spotted onto YPD plates and an incubated at 37°C for 3 days.

### Murine infection model

*C. neoformans* inoculum was prepared by culturing cells in 5 mL YPD on a tissue culture roller drum at 30°C for approximately 16 hours. Cells were collected by centrifugation, washed twice with sterile phosphate-buffered saline (PBS), and the cell density was determined with a hemocytometer. The final cell concentration was adjusted to 4 x 10^7^ /mL in PBS. Four-to five week-old A/J mice (Jackson Laboratory, USA) were utilized for the murine intranasal infection model (*n*=14 for each group, 7 male and 7 female). Mice were anesthetized with isoflurane and infected by intranasal instillation of 25 μL inoculum (10^6^ cells). Mice survival was monitored daily, and euthanasia was performed via CO_2_ exposure upon reaching humane endpoints, including greater than 20% weight loss, reduced grooming and mobility, or a hunched appearance. For fungal burden analysis, four mice (2 male, 2 female) from each group were randomly selected and euthanized via CO_2_ exposure 14 days post-infection. The brain and lungs were dissected and homogenized in 800 μL sterile PBS using bead-beating. Organ homogenates were plated onto YPD agar containing antibiotics (100 μg/mL ampicillin, 100 μg/mL kanamycin) to isolate fungal colonies. Survival data were plotted using Kaplan-Meier curves and statistically analyzed through log-rank (Mantel-Cox) test. Statistical analyses of fungal burdens were performed using either Mann-Whitney U test or one-way ANOVA with Dunnett’s multiple comparisons test. Data plotting and analysis of mouse survival and fungal burden was performed with GraphPad Prism v 10.2.3.

### Analysis of RNA-seq and Ribo-seq data

Mating crosses were performed in Murashige and Skoog (MS) medium and checked for filamentation under dissecting microscope. On day 7, mating filaments were harvested by scraping and flash frozen in liquid N_2._ Frozen cell pellets were lyophilized overnight and pulverized for 30 seconds in the bead beater with sterile zirconium beads (0.5 mm diameter). RNA extraction was performed as per instructions of the PureLink RNA Mini Kit from Ambion. Corall total RNA-seq library preparation kit from Lexogen was used as per manufacturer instructions to generate the RNA-seq library. Ribosomal profiling workflow for *C. neoformans* mating samples was modified from published methods from Ingolia lab (50). Sequencing of the strains was performed at the Duke sequencing facility core (https://genome.duke.edu/).

RNA-seq reads were mapped with Hisat2 v2.2.1 (51) to the H99 genome(52). Reads mapping to annotated features were counted as described (53) with the modification that reads were strand-specific and were only counted if they mapped to the strand of the corresponding feature. Ribo-seq reads were trimmed with cutadapt (v3.4) (54) with parameters –j 16 –e 0.1 –O 4 –a AGATCGGAAGAGCACACGTCTGAAC –m 25 –max-n 0. Trimmed reads were mapped to the *C. neoformans* rRNA and tRNA loci using bowtie2 (v2.4.4) (55), and only reads that did not map to these loci were used for downstream analysis. Reads were demultiplexed based on adapter sequences using cutadapt and mapped with STAR (v2.7.8a) (56) to the H99 genome. Reads mapping to annotated features were counted in Bioconductor (57) (in R v4.1 using Rstudio 2021.09.1 (58)) based on GenomicAlignments and GenomicFeatures (59). To analyze sample distances, read counts for RNA-seq and Ribo-seq data were analyzed in R (v4.1.2) (60) with DESseq2 (v1.34) (61). To analyze gene regulation levels, RNA-seq and Ribo-seq read counts were analyzed in R (v4.1.2) with anota2seq (v1.16.0) (30). Parameters for differential gene expression for anota2seq were a maxPAdj=0.05 and minEff=1.

## Acknowledgments

We thank Anna Floyd-Averette for invaluable assistance with the mouse experiments and constant support. We also thank all the members of the Heitman Lab for constructive suggestions, Dr. Blake R. Billmyre (University of Georgia) for providing and communication of the essentiality prediction of *PRT1*, Dr. Leah E. Cowen and Dr. Ci Fu (University of Toronto) or providing the diploid strain CnLC6683. This study was supported by NIH/NIAID R01 grants AI039115-27, AI050113-20, and AI133654-07 awarded to J.H. T.A.D. and U.K. are funded by the German Research Foundation (DFG; Bonn Bad Godesberg, Germany; grant no. KU 517/15-1). M.N. is funded by the DFG (grant NO407/7-2). M.S.S is funded by the R21 (AI138158). Dr. Joseph Heitman is codirector and fellow of the Canadian Institute for Advanced Research (CIFAR) program Fungal Kingdom: Threats & Opportunities.

## Data Availability Statement

Raw data are available at Bioproject: PRJNA1191513.

## Notes

### Competing Interest Statement

The authors have declared no competing interest.

