## supplemental material for "Essential genes encoded by the mating-type locus of the human fungal pathogen *Cryptococcus neoformans*"

**SUPPLEMENTAL TABLES AND FIGURES**

**TableS1. Strains used in this study.**

| **Strain** | **Background strain** | **Genotype** | **Note** |
| --- | --- | --- | --- |
| H99α |  | WT | (1) |
| KN99**a** |  | WT | (1) |
| KN99α |  | WT | (1) |
| CnLC6683 |  | WT | (2) |
| ZB225 | CnLC6683 | *myo2***a**Δ::*NAT*/*MYO2*α | This study |
| ZB505 | CnLC6683 | *myo2***a**Δ::*NAT*/*MYO2*α | This study |
| ZB226 | CnLC6683 | *prt1***a**Δ::*NAT* /*PRT1*α | This study |
| ZB228 | CnLC6683 | *prt1***a**Δ::*NAT* /*PRT1*α | This study |
| ZB229 | CnLC6683 | *rpl22***a**Δ::*NAT* /*RPL22*α | This study |
| ZB507 | CnLC6683 | *rpl22***a**Δ::*NAT* /*RPL22*α | This study |
| ZB231 | CnLC6683 | *rpl39***a**Δ::*NAT* /*RPL39*α | This study |
| ZB511 | CnLC6683 | *rpl39***a**Δ::*NAT* /*RPL39*α | This study |
| ZB314 | CnLC6683 | *rpo41***a**Δ::*NAT* /*RPO41*α | This study |
| ZB315 | CnLC6683 | *rpo41***a**Δ::*NAT* /*RPO41*α | This study |
| ZB237 | CnLC6683 | *MYO2***a/***myo2*αΔ::*NEO* | This study |
| ZB318 | CnLC6683 | *PRT1***a/***prt1*αΔ::*NEO* | This study |
| ZB512 | CnLC6683 | *PRT1***a/***prt1*αΔ::*NEO* | This study |
| ZB238 | CnLC6683 | *RPL22***a/***rpl22*αΔ::*NEO* | This study |
| ZB583 | CnLC6683 | *RPL22***a/***rpl22*αΔ::*NEO* | This study |
| ZB232 | CnLC6683 | *RPL39***a/***rpl39*αΔ::*NEO* | This study |
| ZB509 | CnLC6683 | *RPL39***a/***rpl39*αΔ::*NEO* | This study |
| ZB234 | CnLC6683 | *RPO41***a/***rpo41*αΔ::*NEO* | This study |
| ZB241 | CnLC6683 | *RPO41***a/***rpo41*αΔ::*NEO* | This study |
| ZB478 | KN99**a** | P_tet_-*MYO2***a** | This study |
| ZB479 | KN99**a** | P_tet_-*MYO2***a** | This study |
| ZB478 | H99α | P_tet_-*MYO2*α | This study |
| ZB479 | H99α | P_tet_-*MYO2*α | This study |
| ZB669 | H99α | P_2x_*_CTR4_*-*TOR1* | (3) |
| ZB670 | KN99**a** | P_2x_*_CTR4_*-*PRT1***a** | This study |
| ZB671 | KN99**a** | P_2x_*_CTR4_*-*PRT1***a** | This study |
| ZB235 | KN99**a** | *myo2***a**Δ::*NAT* | This study |
| ZB236 | KN99**a** | *myo2***a**Δ::*NAT* | This study |
| ZB242 | KN99α | *myo2*αΔ::*NEO* | This study |
| ZB243 | KN99α | *myo2*αΔ::*NEO* | This study |
| ZB487 | KN99**a**, recombinant mitochondria | Hem15-GFP--*NAT* Nop1-mCherry--*NEO* | This study |
| ZB307 | KN99α, recombinant mitochondria | *myo2*Δ::*NEO* | This study |
| ZB307 | KN99α, recombinant mitochondria | *myo2*Δ::*NEO* | This study |

**Table S2. Primers used in this study.**

| **Name** | **Squence 5'-3'** | **Purpose** |
| --- | --- | --- |
| JOHE45803/SS | CTCCACCCTTACTGATCGCC | *MYO2*_**a**_Upstream_Forward |
| JOHE45804/SS | GTCATAGCTGTTTCCTGAGTACCAGGAACCCAGCTAGT | *MYO2*_**a**_Upstream_Reverse_M13 |
| JOHE45805/SS | CTGGCCGTCGTTTTACTATGCGTGCAGATGTACCGG | *MYO2*_**a**_Downstream_Forward_M13 |
| JOHE45806/SS | ACTCCTTCCTCGGCTCTTCT | *MYO2*_**a**_Downstream_Reverse |
| JOHE45807/SS | CGAGAATGAAGAGGGCTGGG | *PRT1*_**a**_Upstream_Forward |
| JOHE45808/SS | GTCATAGCTGTTTCCTGCACTGACTTTTCCTCGATGTCC | *PRT1*_**a**_Upstream_Reverse_M13 |
| JOHE45809/SS | CTGGCCGTCGTTTTACCAGAGAGATCGGGGAGTGGA | *PRT1*_**a**_Downstream_Forward_M13 |
| JOHE45810/SS | ACACAATCAATGCGCCTGC | *PRT1*_**a**_Downstream_Reverse |
| JOHE45811/SS | ACCTTCACCTCAAGCTCTGC | *RPL22*_**a**_Upstream_Forward |
| JOHE45812/SS | GTCATAGCTGTTTCCTGGCAGCATCCACAACCCAAAA | *RPL22*_**a**_Upstream_Reverse_M13 |
| JOHE45813/SS | CTGGCCGTCGTTTTACGTGTCGTTGCCACTTCCAAG | *RPL22*_**a**_Downstream_Forward_M13 |
| JOHE45814/SS | TGATGTCGGCAACCTTCCTC | *RPL22*_**a**_Downstream_Reverse |
| JOHE45815/SS | TTTCCATCCTCGTCGCTGTT | *RPL39*_**a**_Upstream_Forward |
| JOHE45816/SS | GTCATAGCTGTTTCCTGGCGTCGAACCAAGCTTAACC | *RPL39*_**a**_Upstream_Reverse_M13 |
| JOHE45817/SS | CTGGCCGTCGTTTTACCAAACAAAGTCTGCCGTGGG | *RPL39*_**a**_Downstream_Forward_M13 |
| JOHE45818/SS | CACTGCATCAATCGAACGCC | *RPL39*_**a**_Downstream_Reverse |
| JOHE45819/SS | GCGCACAACACTCCATTTAGAC | *RPO41*_**a**_Upstream_Forward |
| JOHE45820/SS | GTCATAGCTGTTTCCTGGATGTCTTGCCGCCTTGTCC | *RPO41*_**a**_Upstream_Reverse_M13 |
| JOHE45821/SS | CTGGCCGTCGTTTTACCCTCTCCCCACTGCTCTACT | *RPO41*_**a**_Downstream_Forward_M13 |
| JOHE45822/SS | GCTTGGGTCTACTATGTCTGTCG | *RPO41*_**a**_Downstream_Reverse |
| JOHE45823/SS | TTCTGAAGTCCCTCGCAACC | *MYO2*_alpha_Upstream_Forward |
| JOHE45824/SS | GTCATAGCTGTTTCCTGGGATCTGTTGTCGCGCTTTG | *MYO2*_alpha_Upstream_Reverse_M13 |
| JOHE45825/SS | CTGGCCGTCGTTTTACCAGACGATGTGGGCCCATAT | *MYO2*_alpha_Downstream_Forward_M13 |
| JOHE45826/SS | TATCTCAGCAGCCACGCAAA | *MYO2*_alpha_Downstream_Reverse |
| JOHE45827/SS | TCGGACACCTCAGCTGTAGA | *PRT1*_alpha_Upstream_Forward |
| JOHE45828/SS | GTCATAGCTGTTTCCTGGCGAGAACAGAGGCAAGTCA | *PRT1*_alpha_Upstream_Reverse_M13 |
| JOHE45829/SS | CTGGCCGTCGTTTTACGCTCAGCTTCAATCTCACGC | *PRT1*_alpha_Downstream_Forward_M13 |
| JOHE45830/SS | GGAACCAGTCATCCCCACTC | *PRT1*_alpha_Downstream_Reverse |
| JOHE45831/SS | CCGGCGTGATGGTAGTGATA | *RPL22*_alpha_Upstream_Forward |
| JOHE45832/SS | GTCATAGCTGTTTCCTGGTAGTGGAGGGAGCTTTCGG | *RPL22*_alpha_Upstream_Reverse_M13 |
| JOHE45833/SS | CTGGCCGTCGTTTTACTTGCCACTTCCAAGGACACC | *RPL22*_alpha_Downstream_Forward_M13 |
| JOHE45834/SS | CCACTTCATCGACGGTAGGG | *RPL22*_alpha_Downstream_Reverse |
| JOHE45847/SS | CGGTATTTCTGAACGCTCCG | *RPL39*_alpha_Upstream_Forward_4 |
| JOHE45848/SS | GTCATAGCTGTTTCCTGGTCGTAGGAGAATAAGGCGGA | *RPL39*_alpha_Upstream_Reverse_4_M13 |
| JOHE45837/SS | CTGGCCGTCGTTTTACTTCCGCCTGAAGACTGACAG | *RPL39*_alpha_Downstream_Forward_M13 |
| JOHE45838/SS | TCGGCCTTGAATGATGACCA | *RPL39*_alpha_Downstream_Reverse |
| JOHE45839/SS | GCTGGGAGTGTCAAAAGCGA | *RPO41*_alpha_Upstream_Forward |
| JOHE45840/SS | GTCATAGCTGTTTCCTGGCAGGAAAGCAGAATTGGGTC | *RPO41*_alpha_Upstream_Reverse_M13 |
| JOHE45841/SS | CTGGCCGTCGTTTTACCCTCTCCCCACTGCTCTACT | *RPO41*_alpha_Downstream_Forward_M13 |
| JOHE45842/SS | TGGGTCTACTATGTCTGTCGA | *RPO41*_alpha_Downstream_Reverse |
| JOHE45853/SS | ATCGATTGGGCCTTCATCTC | *MYO2*_**a**_Internal_Forward |
| JOHE45854/SS | AGGAATCCATGGCCGCATTG | *MYO2*_**a**_Internal_Reverse |
| JOHE45855/SS | AATCAGCTGGAAATTCATTG | *MYO2*_alpha_Internal_Forward |
| JOHE45856/SS | CTGGGATGGAGCTTCTGG | *MYO2*_alpha_Internal_Reverse |
| JOHE45857/SS | CCAATCCAGAAAGAGATGGC | *PRT1*_**a**_Internal_Forward |
| JOHE45858/SS | GGTCCAACGCCATTACATTAG | *PRT1*_**a**_Internal_Reverse |
| JOHE45859/SS | AATCATCAATCTAGTTGCCA | *PRT1*_alpha_Internal_Forward |
| JOHE45860/SS | TTGATATCATGCCAATAATGAC | *PRT1*_alpha_Internal_Reverse |
| JOHE45861/SS | TCGCCGCTTTTGAGAAGTTT | *RPL22*_**a**_Internal_Forward |
| JOHE45862/SS | TCTTGGTAAGGTACTTAAGA | *RPL22*_**a**_Internal_Reverse |
| JOHE45863/SS | TTGCCGCGTTTGAGAAGTTC | *RPL22*_alpha_Internal_Forward |
| JOHE45864/SS | TCTTCGTAAGGTACTTGAGG | *RPL22*_alpha_Internal_Reverse |
| JOHE45865/SS | TCACGATATCGACAACACCGC | *RPO41*_**a**_Internal_Forward |
| JOHE45866/SS | TTTGACGTTGTGCTCGAGTC | *RPO41*_**a**_Internal_Reverse |
| JOHE45867/SS | TCACGATATCAACGACACCGG | *RPO41*_alpha_Internal_Forward |
| JOHE45866/SS | TTTGACGTTGTGCTCGAGTC | *RPO41*_alpha_Internal_Reverse |
| JOHE45868/SS | GGTTAAGCTTGGTTCGACGCCA | *RPL39*_**a**_Internal_Forward |
| JOHE45869/SS | ATGCTGACTCCTGTCCCAAATT | *RPL39*_**a**_Internal_Reverse |
| JOHE45870/SS | GGTTGAGCTTGGTTCGACGCCA | *RPL39*_alpha_Internal_Forward |
| JOHE45871/SS | ATGCTGACGTCTGCCCTCAATC | *RPL39*_alpha_Internal_Reverse |
| JOHE45876/SS | AGATCGATGGTGGGGAGGAA | *MYO2*_**a**_Junction_Forward |
| JOHE45877/SS | CCAAGATGTTTACGTTCGGG | *MYO2*_**a**_Junction_Reverse |
| JOHE45878/SS | GGAACCAGTCATCCCCACTC | *MYO2*_alpha_Junction_Forward |
| JOHE45879/SS | GCAAAGGACCCATCTCAGCT | *MYO2*_alpha_Junction_Reverse |
| JOHE45880/SS | CGAGACGACTGGAATGGTGT | *PRT1*_**a**_Junction_Forward |
| JOHE45881/SS | AGCCATGTGTGAATCCTGCG | *PRT1*_**a**_Junction_Reverse |
| JOHE45884/SS | AGAACTGTGCCCGGAATAGC | *PRT1*_alpha_Junction_Forward2 |
| JOHE45885/SS | AGGGCTTGCCGAAGAACAAT | *PRT1*_alpha_Junction_Reverse2 |
| JOHE45888/SS | GACCAATGCCGGAAGAGGAT | *RPL22*_**a**_Junction_Forward |
| JOHE45889/SS | GTGAGTACCGCATTACCAGC | *RPL22*_**a**_Junction_Reverse |
| JOHE45890/SS | CTTCAGCCTCAGACTCACCC | *RPL22*_alpha_Junction_Forward |
| JOHE45891/SS | CCATTGTCCATGTTCCCCGT | *RPL22*_alpha_Junction_Reverse |
| JOHE45892/SS | CCAAGTCTCTGCTTCCACCA | *RPL39*_**a**_Junction_Forward |
| JOHE45893/SS | TGAGGACAGATTGGCGTGAG | *RPL39*_**a**_Junction_Reverse |
| JOHE45894/SS | CCCTTTCCATCACCTCCGAT | *RPL39*_alpha_Junction_Forward |
| JOHE45895/SS | GGGTTGAAGCTGGGGAGAAC | *RPL39*_alpha_Junction_Reverse |
| JOHE45896/SS | GGTTCGGGCGCTAAGTAACA | *RPO41*_**a**_Junction_Forward |
| JOHE45897/SS | ATCTCGCCGCAAATACCACT | *RPO41*_**a**_Junction_Reverse |
| JOHE45898/SS | TTTAGGCATGGACGCACAGT | *RPO41*_alpha_Junction_Forward |
| JOHE45899/SS | ATCTCGCCGCAAATACCACT | *RPO41*_alpha_Junction_Reverse |
| JOHE52755ZB26 | ACCGGCAGGGTATACTGTTGGCGCTTTGTAAGGTGGACAAGTTTTAGAGCTAGAAATAGC | *MYO2*_alpha_5'_gRNA |
| JOHE52756ZB27 | ACCGGCAGGGTATACTGTTGGCTTGGTAGACGTCCCAGCGGTTTTAGAGCTAGAAATAGC | *MYO2*_alpha_3'_gRNA |
| JOHE52757ZB28 | ACCGGCAGGGTATACTGTTGATGTCCGCCCCCTACAGAAAGTTTTAGAGCTAGAAATAGC | *MYO2*_**a**_5'_gRNA |
| JOHE52758ZB29 | ACCGGCAGGGTATACTGTTGCTGCCAATCGCCGGACAGATGTTTTAGAGCTAGAAATAGC | *MYO2*_**a**_3'_gRNA |
| JOHE52890/ZB109 | ACCGGCAGGGTATACTGTTGTATGCTGTGACCGCTCAGAAGTTTTAGAGCTAGAAATAGC | *PRT1*_**a**_5'_gRNA |
| JOHE52891/ZB110 | ACCGGCAGGGTATACTGTTGCTGGGCACTGGGGAACATTTGTTTTAGAGCTAGAAATAGC | *PRT1*_**a**_3'_gRNA |
| JOHE52892/ZB111 | ACCGGCAGGGTATACTGTTGGCAGCAGCTGGTACTAACAAGTTTTAGAGCTAGAAATAGC | *PRT1*_alpha_5'_gRNA |
| JOHE52893/ZB112 | ACCGGCAGGGTATACTGTTGTCCTCCAACGCTGCTTAGTAGTTTTAGAGCTAGAAATAGC | *PRT1*_alpha_3'_gRNA |
| JOHE52760ZB31 | ACCGGCAGGGTATACTGTTGCAAGTACTTCAAGGTTGATCGTTTTAGAGCTAGAAATAGC | *RPL22*_alpha_3'_gRNA_GI |
| JOHE52761ZB32 | ACCGGCAGGGTATACTGTTGGTTTGCAATCAGGATTACATGTTTTAGAGCTAGAAATAGC | *RPL22*_alpha_5'_gRNA |
| JOHE52763ZB34 | ACCGGCAGGGTATACTGTTGAGTTGCAATGAGGGTTACATGTTTTAGAGCTAGAAATAGC | *RPL22*_**a**_5'_gRNA_GI |
| JOHE52765ZB36 | ACCGGCAGGGTATACTGTTGTGTATCTACACGTTACTAACGTTTTAGAGCTAGAAATAGC | *RPL22*_**a**_gRNA3_GI |
| JOHE52894/ZB113 | ACCGGCAGGGTATACTGTTGCCACGGCAGACTTTGTTTGCGTTTTAGAGCTAGAAATAGC | *RPL39*_**a**_5'_gRNA |
| JOHE52895/ZB114 | ACCGGCAGGGTATACTGTTGATAACGCCAAGCGTCGTCATGTTTTAGAGCTAGAAATAGC | *RPL39*_**a**_3'_gRNA |
| JOHE52896/ZB115 | ACCGGCAGGGTATACTGTTGGGAGTCGTAGGAGAATAAGGGTTTTAGAGCTAGAAATAGC | *RPL39*_alpha_5'_gRNA |
| JOHE52897/ZB116 | ACCGGCAGGGTATACTGTTGAGGCGGAACCACTGAGGAAGGTTTTAGAGCTAGAAATAGC | *RPL39*_alpha_3'_gRNA |
| JOHE52898/ZB117 | ACCGGCAGGGTATACTGTTGGAGCACTAGACCCATGGTTGGTTTTAGAGCTAGAAATAGC | *RPO41*_**a**_5'_gRNA |
| JOHE52899/ZB118 | ACCGGCAGGGTATACTGTTGGAGGATTTCCTTGACCGGTAGTTTTAGAGCTAGAAATAGC | *RPO41*_**a**_3'_gRNA |
| JOHE52900/ZB119 | ACCGGCAGGGTATACTGTTGGTGACAGCGGCTTTGGATTCGTTTTAGAGCTAGAAATAGC | *RPO41*_alpha_5'_gRNA |
| JOHE52901/ZB120 | ACCGGCAGGGTATACTGTTGTTTCTACGATCTTATCGGCAGTTTTAGAGCTAGAAATAGC | *RPO41*_alpha_3'_gRNA |
| JOHE52738ZB9 | TGTAAAACGACGGCCAGT | M13F |
| JOHE52739ZB10 | CAGGAAACAGCTATGAC | M13R |
| JOHE52740ZB11 | CATGCATCTAGGTCTAGAAACC | Cas9-F |
| JOHE52741ZB12 | CCTCTTCACGTGGACGCTCC | Cas9-R |
| JOHE52742ZB13 | GCCCTAGTCCATTGCGAACG | U6-F |
| JOHE52743ZB14 | CAACAGTATACCCTGCCGGTG | U6-R |
| JOHE52744ZB15 | GTTTTAGAGCTAGAAATAGCAAG | sgRNA-F |
| JOHE52745ZB16 | AAGATACTCGATTTGCCGTCC | sgRNA-R |
| JOHE52746ZB17 | GCTCATGGATCCTTTGCATTAGAACTAAAAACAAAGCA | sgRNA-final-F |
| JOHE52747ZB18 | GATCATCCGCGGTAAAACAAAAAAGCACCGACTCGGTGCC | sgRNA-final-R |
| JOHE54481/YC183 | TAGGCCCCTTTTCCGTCTAT | *PRT1***a** *CTR4* promoter replacement L1 |
| JOHE54482/YC184 | CACTCGAATCCTGCATGCTTTTGCGTATGGCGTGGTG | *PRT1***a** *CTR4* promoter replacement L2 |
| JOHE54483/YC185 | CGACAACGACTTCACCAATCATGTCGGTCACCGACTTAAC | *PRT1***a** *CTR4* promoter replacement R1 |
| JOHE54484/YC186 | GCCATCTCTTTCTGGATTGG | *PRT1***a** *CTR4* promoter replacement R2 |
| JOHE54314/ZB363 | GCATGCAGGATTCGAGTG | NAT/CTR-L |
| JOHE54315/ZB364 | GATTGGTGAAGTCGTTGTCG | NAT/CTR-R |
| JOHE54314/ZB366 | AAGGTGTTCCCCGACGACGAATCG | NAT-SM1 |
| JOHE54315/ZB367 | CGATTCGTCGTCGGGGAACACCTT | NAT-SM2 |
| JOHE53219/ZB245 | ACCGGCAGGGTATACTGTTGCTGACTTAGTTAGCTAAGATGTTTTAGAGCTAGAAATAGC | *PRT1***a**_dox_gRNA_1 |
| JOHE53194/ZB220 | ATCGCTGTATGTCTCCGAATCG | *PRT1***a**_dox_internal-R |
| JOHE53195/ZB221 | ATACTTGTTGAAACGGGCTATG | *PRT1***a**_dox_internal-F |
| JOHE54719/YC99 | TGTGGATGCTGGCGGAGGATA | B79 5' screening primer, Screening oligo on *ACT* promoter |
| JOHE54719/YC100 | TTCCCACCCTCAGCAACGCC | J12579 3' screening primer, Screening oligo on *TRP* terminator |
| JOHE53175/ZB201 | TAACAACGGGGTCCAGAAATCG | *MYO2***a**_dox_Upstream-F |
| JOHE53176/ZB202 | CACTGGCCGTCGTTTTACAAGAGTATGTGAGATGAGTGATGC | *MYO2***a**_dox_Upstream-R |
| JOHE53177/ZB203 | CATGGTCATAGCTGTTTCCTATGTCCGCCCCCTACAGAAAA | *MYO2***a**_dox_Downstream-F |
| JOHE53178/ZB204 | AGGAGTAAACGGATTAACCGAC | *MYO2***a**_dox_Downstream-R |
| JOHE53179/ZB205 | CGTCTTTACGTCTCATTATTGG | *MYO2***a**_dox_Internal-F |
| JOHE53180/ZB206 | GTACTGGTAGCGAACCAATTT | *MYO2***a**_dox_Internal-R |
| JOHE53181/ZB207 | CGTCGGGAGGTCGATTTTTCTA | *MYO2***a**_dox_Junction-F |
| JOHE53182/ZB208 | CCCTTCTTCTGACCCGAGTATA | *MYO2***a**_dox_Junction-R |
| JOHE53183/ZB209 | AGCTGTGTTCCTTACGCTGCAA | *MYO2*α_dox_Upstream-F |
| JOHE53184/ZB210 | CACTGGCCGTCGTTTTACAACAGGCCCTTGTGACAAGATTCG | *MYO2*α_dox_Upstream-R |
| JOHE53185/ZB211 | CATGGTCATAGCTGTTTCCTATGACTTCTTTATATTCAAAGG | *MYO2*α_dox_Downstream-F |
| JOHE53186/ZB212 | GTTGAGTGGAGTAAACGGATTC | *MYO2*α_dox_Downstream-R |
| JOHE53187/ZB213 | ATACCTTATAGACTCAGTAGCG | *MYO2*α_dox_Internal-F |
| JOHE53188/ZB214 | ACGAGGACGTCTAATGGGTTG | *MYO2*α_dox_Internal-R |
| JOHE53189/ZB215 | CATGACACTGATGTACATCGCC | *MYO2*α_dox_Junction-F |
| JOHE53190/ZB216 | CCTTCTTTTGACCTGCGTACAG | *MYO2*α_dox_Junction-R |
| JOHE53215/ZB241 | ACCGGCAGGGTATACTGTTGGAACAGTCTTCTGAAAATTGGTTTTAGAGCTAGAAATAGC | *MYO2***a**_dox_gRNA |
| JOHE53217/ZB243 | ACCGGCAGGGTATACTGTTGTCTAAATATCATCCGGCAAGGTTTTAGAGCTAGAAATAGC | *MYO2*α_dox_gRNA |

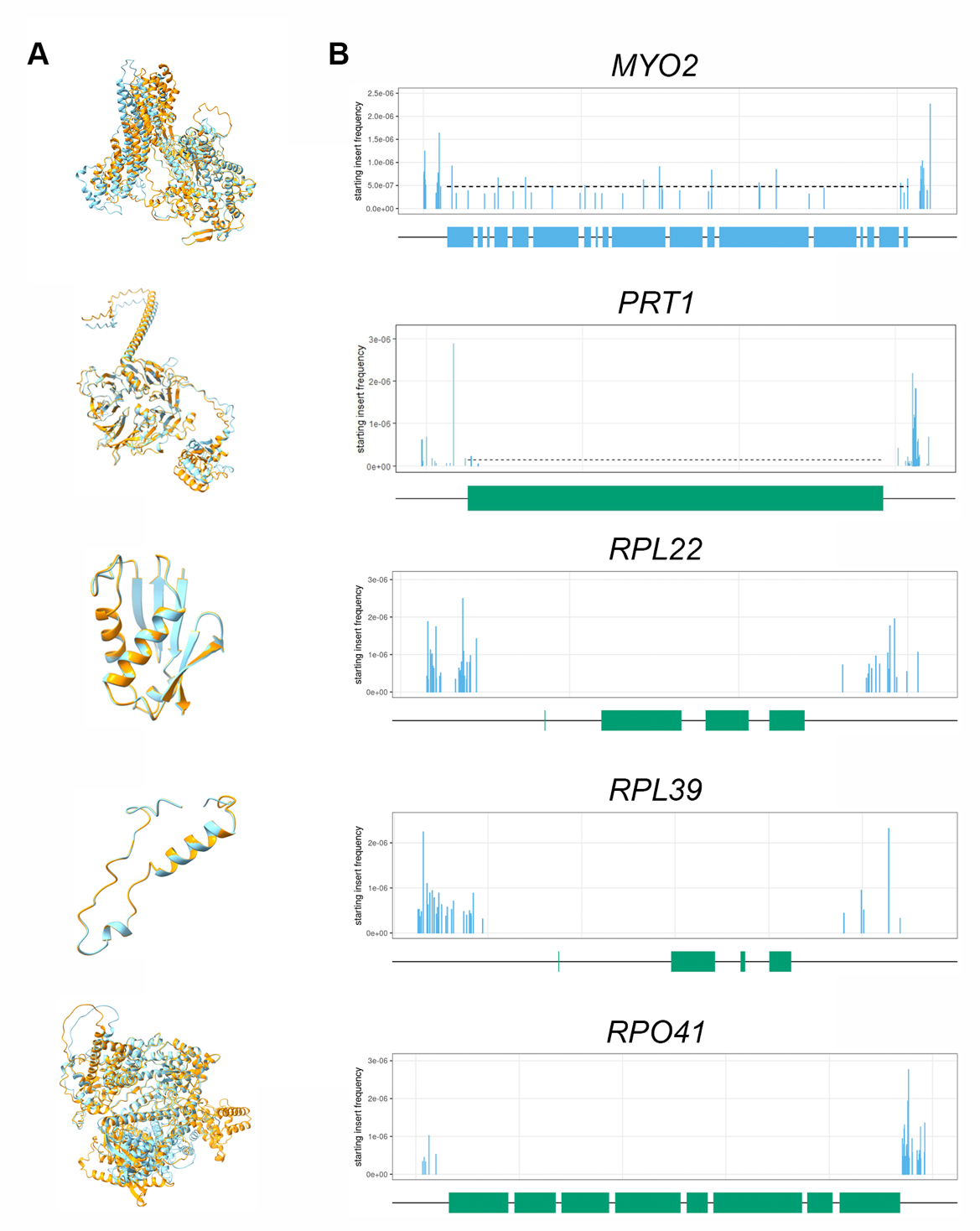

**FIG S1.** Predicted protein structures and essentiality for *MYO2*, *PRT1*, *RPL22*, *RPL39*, and *RPO41*. (A) Protein structures encoded by the **a** and α alleles for each gene were predicted using AlphaFold and aligned with the program ChimeraX, with **a** alleles depicted in orange and α alleles in blue. (B) Predicted essentiality of the α allele of each gene based on Tn-seq analyses (https://simrcompbio.shinyapps.io/Crypto_TN_seq_viewer/).

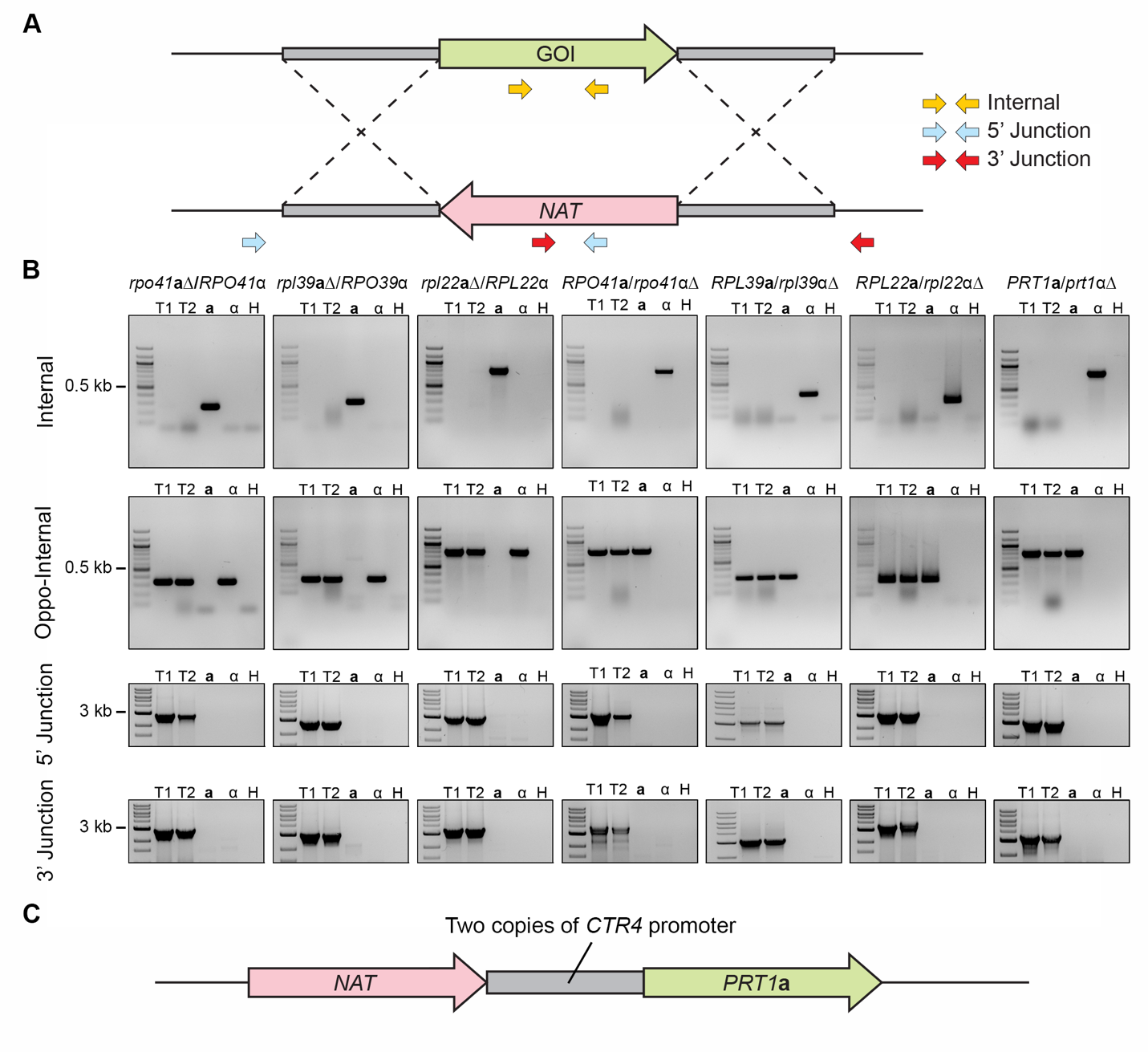

**FIG S2.** Genotypic validation of heterozygous deletion mutants. (A) Diagram of the gene replacement approach utilized to generate heterozygous deletion mutants and positions of primer pairs utilized for genotypic validation of transformants. (B) Genotyping of the heterozygous deletion mutants was conducted with primers targeting the internal regions of the ORFs of deleted alleles and alleles of the opposite mating type, as well as the 5’- and 3’-junctions of the deletion alleles; **a**, α, and H indicate the KN99**a**, KN99α, and water controls for PCR, respectively. See Fig. 2A and Fig. S5C for validation PCR of *myo2***a**Δ/*MYO2*α and *MYO2***a**/*myo2*αΔ, respectively. (C) Diagram of the tandem *CTR4* promoter insertion approach utilized to generate mutants for *PRT1***a**.

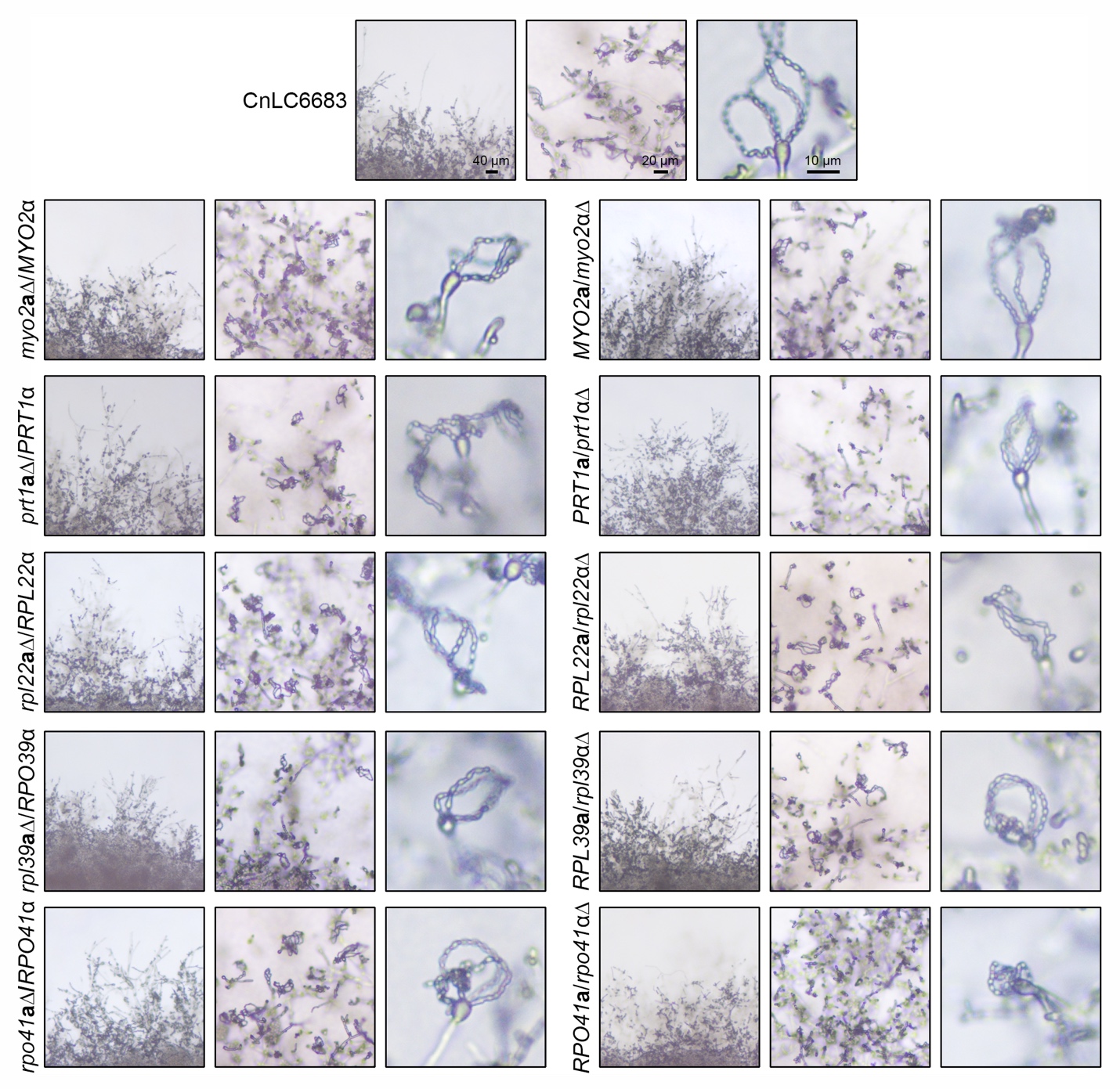

**FIG S3.** Selfing of heterozygous deletion mutants on MS plates. Light microscopy images showing robust sporulation in all mutants except *PRT1***a**/*prt1*αΔ and *RPO41***a**/*rpo41*αΔ. Selfing of *PRT1***a**/*prt1*αΔ and *RPO41***a**/*rpo41*αΔ exhibited robust hyphal development but infrequent sporulation events. Scale bar is indicated in images of CnLC6683 samples. Scale bar = 40 μm (left), 20 μm (middle) and 10 μm (right)

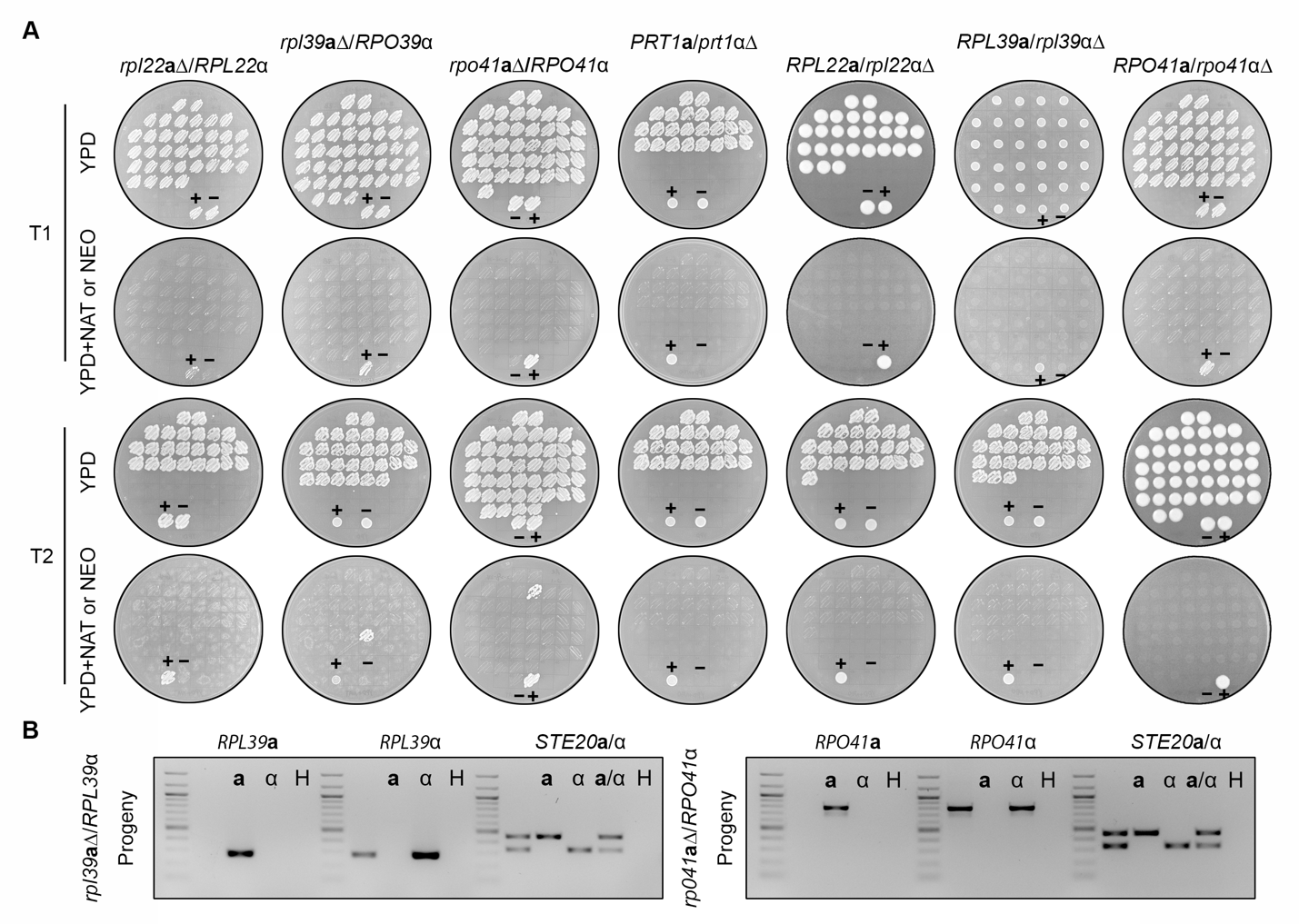

**FIG S4.** Phenotypic analyses of the random spores dissected from heterozygous deletion mutants. (A) Growth of the progeny on solid YPD and YPD supplemented with NAT (deletion of **a** alleles) or NEO (deletion of α alleles). The control (lower) patches are corresponding heterozygous mutants as positive control (+) and wild-type strain CnLC6683 as negative control (-). (B) The two drug resistant progeny from *rpl39***a**Δ/*RPL39*α and *rpo41***a**Δ/*RPO41*α were confirmed to still possess the wildtype allele of the opposite mating type, indicating these resistant progeny are aneuploid for the mating type locus chromosome.

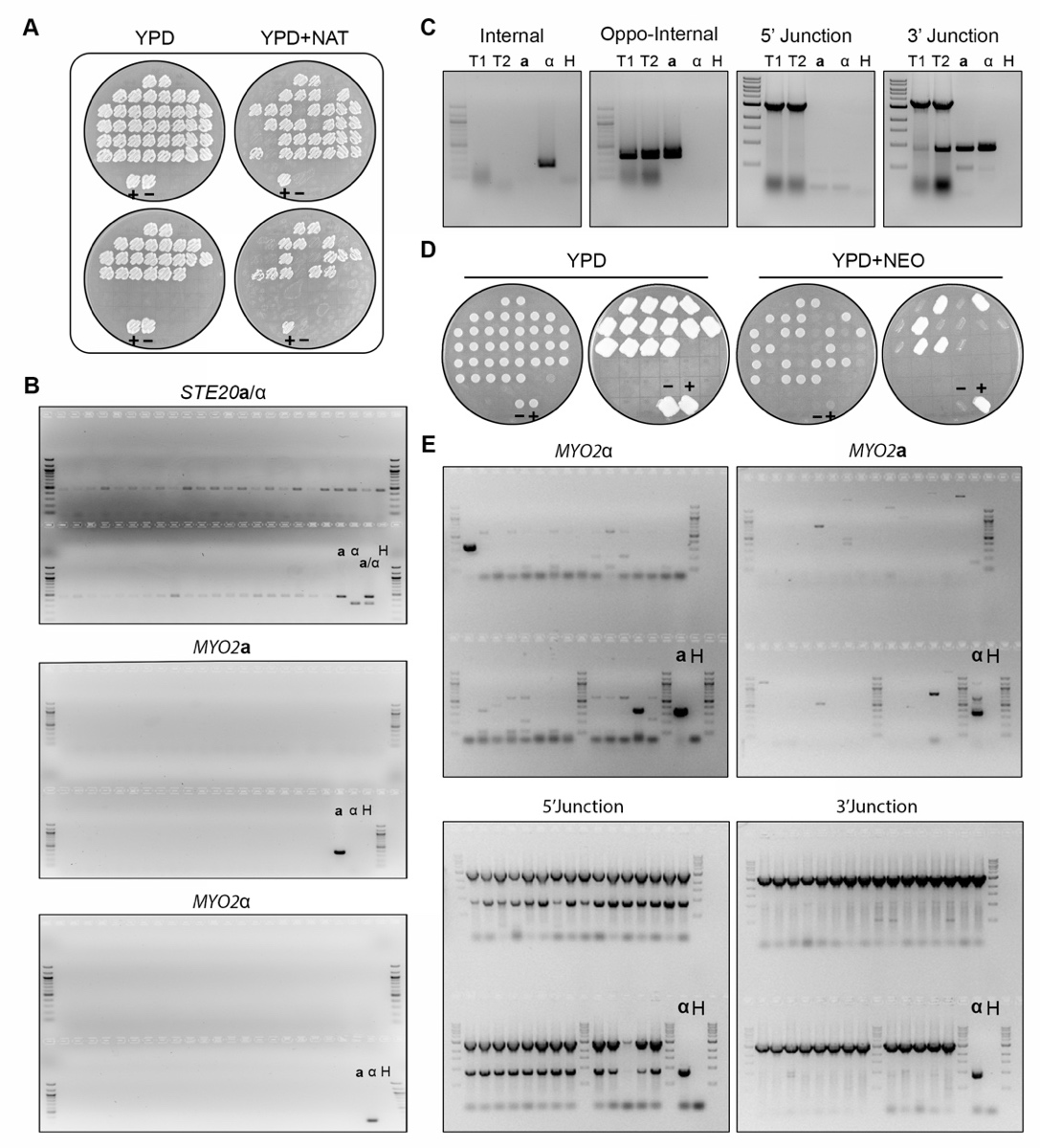

**FIG S5.** *MYO2***a** and *MYO2*α are not essential. (A) Phenotyping of randomly dissected spores of independent *myo2***a**Δ**/***MYO2*α mutant on YPD and YPD+NAT plates. The control (lower) patches are corresponding heterozygous mutants as positive control (+) and wild-type strain CnLC6683 as negative control (-). (B) Random spores dissected from *myo2***a**Δ**/***MYO2*α heterozygous mutants for mating type were genotyped to show the absence of both *MYO2***a** and *MYO2*α. (C) The *MYO2***a/***myo2*αΔ heterozygous mutants were genotypically validated. (D) Random spores dissected from the two independent *MYO2***a/***myo2*αΔ mutants were phenotyped on solid YPD and YPD+NEO plates. Bottom two patches are corresponding heterozygous mutants as positive control (+) and wild-type strain CnLC6683 as negative control (-). (E) Genotyping of *MYO2***a/***myo2*αΔ spores for the absence of both *MYO2***a** and *MYO2*α alleles is shown.

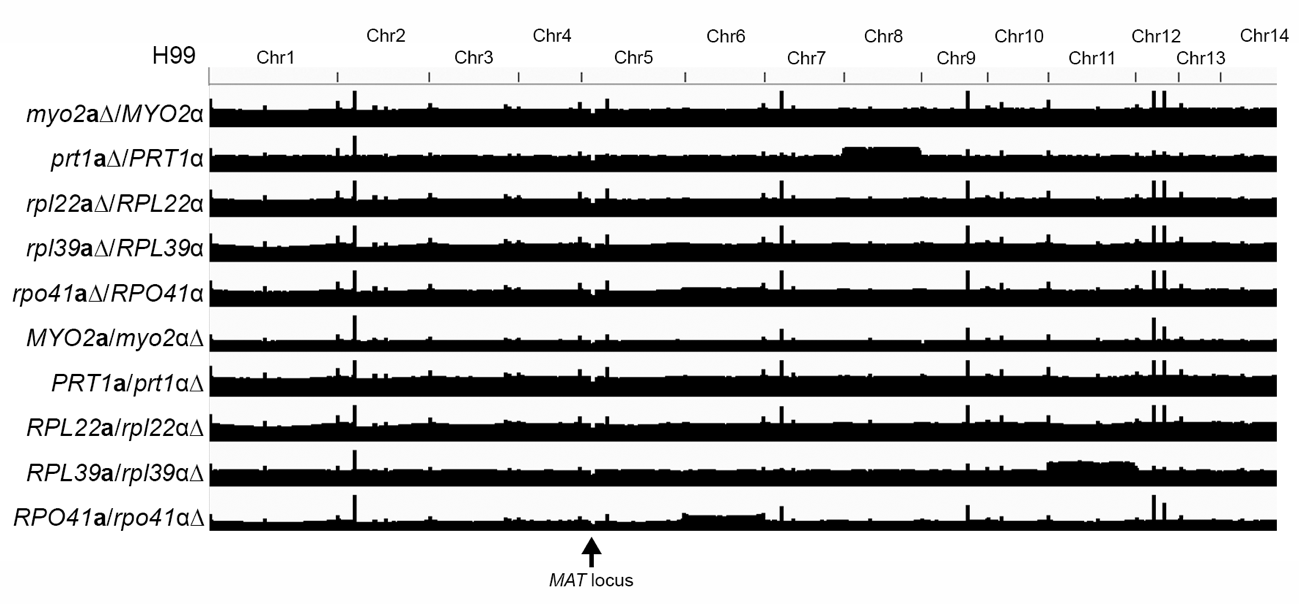

**FIG S6.** Genome-wide read depth analyses of heterozygous deletion mutants with Illumina whole genome sequencing reads. While mutant strains exhibited occasional aneuploidy for some chromosomes, there was no segmental deletion that was inside or linked to the mating type locus, confirming that the inviability observed in the meiotic progeny was due to the absence of the gene that was deleted.
